# The plant specific Histone Lysine Demethylase MtPKDM9B mediates the root nodule symbiosis by controlling H3K27me3 levels and expression of symbiotic genes

**DOI:** 10.64898/2026.09.12.750751

**Authors:** Milagros Ferrari, Milagros Yacullo, Soledad Traubenik, Ezequiel Ibarra, Flavio A Blanco, Mauricio Reynoso, María Eugenia Zanetti

**Affiliations:** Instituto de Biotecnología y Biología Molecular, Facultad de Ciencias Exactas, Universidad Nacional de La Plata, Centro Científico y Tecnológico-La Plata, Consejo Nacional de Investigaciones Científicas y Técnicas, La Plata (CP1900), Argentina

**Author notes:** Institute of Plant Sciences Paris-Saclay (IPS2), CNRS, INRA, Universities Paris-Sud, Evry and Paris-Diderot, Sorbonne Paris-Cite, University of Paris-Saclay, Batiment 630, 91405 Orsay, France.

**Keywords:** alternative splicing, histones posttranslational modifications, nodulation, nitrogen fixing symbiosis, translation

## Abstract

Under nitrogen limiting conditions, legume plants interact with nitrogen fixing bacteria known as rhizobia, resulting in the formation of a new organ, the nodule. This process is accompanied by dramatic changes in gene expression, which operate at different levels. A previous study revealed that histone methylation is differentially modulated during nodulation. However, the histone methyl transferases and demethylases involved in this modulation have not been characterized. In this study we report the identification of the *Medicago truncatula* putative histone lysine demethylase *MtPKDM9B*, which is subject to alternative splicing (AS), and the differential modulation of AS variants at translational level during nodule symbiosis. Knockdown of *MtPKDM9B* impaired infection by rhizobia, nodule development, bacterial viability and the expression of the leghemoglobin coding gene *MtLHB1*. *MtPKDM9B* is the putative ortholog of Arabidopsis *EARLY FLOWERING 6* (*ELF6/AtPKDM9B)* gene involved in the removal of the repressive mark H2K27me3. A combination of ChIP-seq and RNA-seq experiments revealed that *MtPKDM9B* is required for demethylation of H3K27me3 in regions nearby or contained within gene bodies of symbiotic genes and the upregulation of the cognate mRNAs in response to rhizobia, including those encoding the putative ubiquitin ligase MtPUB2, the MYB transcription factor MtMYB040 and the auxin conjugating enzyme MtGH3 (Gretchen Hagen 3). Our findings illustrate how AS and translational regulation of this plant specific histone lysine demethylase contributes to the removal of the repressive mark H3K27me3, promoting transcriptional activation of symbiotic genes required for the formation of functional nitrogen fixing nodules.

## Introduction

Histone post-translational modifications play essential roles in the epigenetic regulation of gene expression in plants shaping developmental programs and responses to environmental cues. One of such modifications is the histone methylation, which can occur in different lysine (K) or arginine (R) residues in the core histone tails. The methylation status of K residues of histone 3 (H3) is highly dynamic and controlled by the activity of histone methyltransferases (writers) and histone demethylases (erasers). Methyl groups in residues of the H3 tail are recognized by specific proteins called readers, which mediate biological functions. In turn, writers, readers and erasers are modulated in response to endogenous and exogenous stimuli, mediating plant growth, development and adaptation to environmental changes (Liu et al., 2010). One, two or three methyl groups can be deposited in different K residues of H3, generating marks that can mediate transcriptional activation (H3K4 and H3K36) or repression (H3K9 and H3K27) at distinct genome locations. Mono-methylation of H3K27 (H3K27me1) is mainly associated with pericentromeric heterochromatin regions, di-methylation of H3K27 (H3K27me2) is found in both euchromatin and facultative heterochromatin, whereas tri-methylation of H3K27 (H3K27me3) is usually associated with transcriptionally repressed genes present in euchromatic regions (Haider and Farrona, 2024).

H3K27 mono- di- and tri- methylation are introduced by the Polycomb repressive complex 2 (PRC2), whose catalytic subunits belong to the group of evolutionarily conserved SET-domain containing proteins (named after three *Drosophila melanogaster* genes: Su[var]3-9, Enhancer of zeste, and Trithorax, which methylate H3K9, H3K27, and H3K4, respectively). In plants, three SET-domain containing proteins, i.e., CURLY LEAF (CLF), SWINGER (SWN) and MEDEA (MEA), are responsible for deposition of H3K27me3 and also contribute to H3K27me2 deposition in euchromatin, whereas H3K27me1 is deposited in heterochromatic regions by the action of ARABIDOPSIS TRITHORAX-RELATED PROTEIN5 (ATXR5) and ATXR6 (Jacob et al., 2010; Xiao et al., 2016). In line with the dynamics of the methylation status, methyl groups can be passively removed due to histone turnover or dilution of histone parental marks, as well as actively removed by proteins with histone lysine demethylase activity (Xiao et al., 2016). Two types of histone lysine demethylases have been identified in plants, animals and fungi: Lysine Specific Demethylase 1 (LSD1) homologues and Jumonji (Jmj) domain-containing proteins, referred to as JUMONJI (JMJ) proteins (Xiao et al., 2016). The JMJ proteins constitute a family that has largely expanded and diverged in angiosperm, presumably due to genome wide duplication events (Qian et al., 2015). The number of gene members varies depending on the species, e.g., 17 in rice (*Oryza sativa*), 19 in maize (*Zea mays*), 21 in Arabidopsis and 27 in poplar (*Populus trichocarpa*). The gene family has been divided into nine subfamilies (Qian et al., 2015). Six of these subfamilies, i.e., lysine demethylases 3 (KDM3), KDM5, Jmj domain 6 (JMJD6) and Putative-Lysine-Specific Demethylases 11, 12 and 13 (PKDM11, PKDM12 and PKDM13) are conserved between plants and vertebrates, whereas the other three subfamilies, PKDM7, PKDM8 and PKDM9, are specific of plants (Qian et al., 2015). In particular, H3K27me3 demethylation has been associated with five JMJ proteins that antagonize the activity of the SET-domain proteins of the PRC2 complex, including EARLY FLOWERING 6 (ELF6, also known as AtPKDM9B), RELATIVE OF EARLY FLOWERING SIX (REF6, also known as AtPKDM9A) (Crevillén et al., 2014; Yan et al., 2018; Antunez-Sanchez et al., 2020), JMJ13 (also known as AtPKDM8) (Yan et al., 2018), JMJ30 (also known as AtPKDM12A) and JMJ32 (also known as AtPKDM13) (Gan et al., 2014).

Dynamic regulation of H3K27me3 marks enables the activation of genes that participate in developmental programs. In consequence, alteration of this dynamic regulation results in severe developmental aberrations. The H3K27me3 demethylases ELF6 and REF6 mediate different biological processes such as flowering, leaf senescence, hormone signaling and control of circadian rhythms (Lu et al., 2011; Crevillén et al., 2014; Gan et al., 2014; Wang et al., 2019). They also play essential functions in the maintenance of genome integrity (Antunez-Sanchez et al., 2020). Although *ELF6* and *REF6* have high sequence identity, they play divergent roles in the control of flowering time in Arabidopsis. *elf6-1* mutants displayed an early floral transition phenotype, whereas three different *ref6* mutants exhibited late flowering under long- and short-day conditions (Noh et al., 2004). It has been also reported that *elf6/ref6* double mutants exhibited pleiotropic phenotypes, including a high number of petals and serrated and downward curled leaves (Yan et al., 2018). *ref6* mutants also exhibit defects in the formation of aerial organ boundaries (Cui et al., 2016). Other JMJ proteins, such as JMJ13, JMJ14, JMJ15, JMJ18, JMJ30, and JMJ32, have been also implicated in the control of flowering (Gan et al., 2014).

H3K27me dynamics also plays a crucial role in the control root architecture (Zanetti et al., 2024). Genes encoding different components of the PRC2 complex, including SWN, EMF2, VRN2, and FERTILIZATION INDEPENDENT ENDOSPERM (FIE), display cell specific expression patterns and function in the control of meristem development and vascular cell proliferation in Arabidopsis roots (de Lucas et al., 2016). More recently, it has been demonstrated that PLETHORA (PLT) transcription factors, which control the patterning of root meristem, interacts with PRC2 and the histone lysine demethylase JMJ703, which removes the active mark H3K4me3, in rice meristems enhancing the H3K27me3/H3K4me3 ratio and maintaining root meristematic genes in a silent state (Li et al., 2026). Moreover, loss of function of *PLT3, PLT4* and *PLT5* exhibited decreased H3K27me3 and enhanced H3K4me3 levels, as well as upregulation of genes harboring these two marks in the root meristem (Li et al., 2026). On the other hand, JMJ14 seems to play functions in conditional root development since mutations in *jmj14* partially suppresses the reduced root meristem size and growth vigor phenotype of *brevis radix* (*brx*) mutants (Cattaneo et al., 2019). Deposition of H3K27me3 repressive mark by the PRC subunits EMF2 or CLF have also shown to inhibit the establishment of founder cells during the initiation of lateral root in Arabidopsis. Moreover, CLF directly binds to the auxin efflux gene *PIN FORMED 1* (*PIN1*) gene, depositing the repressive mark H3K27me3, down-regulating auxin maxima in these cells and thus preventing lateral formation (Gu et al., 2014).

In a previous study we analyzed alternative spliced (AS) transcript variants differentially regulated at the translational level during early stages of the root nodule symbiosis between *Medicago truncatula* and its partner *Sinorhizobium meliloti* (Traubenik et al., 2020). This symbiotic interaction is initiated by the exchange of signals between both organisms, leading to the formation of a new post-embryonic organ called the nodule, where bacteria allocate and fix atmospheric nitrogen using photosynthetic carbohydrates from the plant (Jhu and Oldroyd, 2023). The formation of nodules requires two highly coordinated morphogenetic programs, the infection by rhizobia and the organogenesis of a new organ that begins with the activation of cell division in the cortex, endodermis and pericycle beneath the site of infection (Jhu and Oldroyd, 2023). The activation of both morphogenetic programs involves a significant change in gene expression that operates at the transcriptional and the translational levels. Our analysis of the translatome (i.e., the population of transcripts associated with ribosomes) revealed that a significant proportion of AS transcript variants modulated at translational level in response to rhizobia encode proteins involved in the deposition and removal of epigenetic marks including a putative lysine specific demethylase belonging to the PKDM9 subfamily, designated here as *M. truncatula* Putative-Lysine-Specific Demethylase 9B (MtPKDM9B). It has been previously shown that the level of the euchromatin repressive mark H3K27me3 in genes contained within symbiotic islands is lower in nodules as compared to roots (Pecrix et al., 2018), suggesting that removal of this epigenetic mark is required for transcriptional activation of symbiotic genes. However, up to date the function of individual demethylases involved in the removal of H3K27me3 during the nitrogen-fixing symbiosis has remained unexplored. Here, we investigated the function of *MtPKDM9B* during the establishment of the root nodule symbiosis. *MtPKDM9B* positively modulates the development of functional nitrogen fixing nodules, although seems to negatively influence primary root growth by controlling cell division in the meristematic and elongation zones of the root when sufficient nitrogen is available. ChIP-seq combined with RNA-seq experiments indicated that MtPKDM9B exerts its influence on nodule development by promoting demethylation and transcriptional activation of key symbiotic genes.

## Results

### *MtPKDM9B* produces two alternative spliced transcript variants differentially modulated during the root nodule symbiosis

A previous study based on the translating ribosome affinity purification technique followed by RNA sequencing (TRAP-seq) identified genes with AS transcript variants differentially modulated at translational level during the root nodule symbiosis in *M. truncatula* (Traubenik et al., 2020). One of these genes, *MtrunA17Chr1g0199511*, encodes a plant specific JMJ histone lysine demethylase. A phylogenetic analysis of the JMJ histone specific demethylases gene families from *M. truncatula* and Arabidopsis revealed that the protein encoded by *MtrunA17Chr1g0199511* clustered in the PKDM9 clade with its counterpart in Arabidopsis AtPKDM9B/ ELF6 (Supplemental Figure 1). The PKDM9 clade also included AtPKDM9A/REF6 and the protein encoded by *MtrunA17Chr3g0116441. MtrunA17Chr1g0199511* is syntenic to ELF6/*AtPKDM9B,* whereas *MtrunA17Chr3g0116441* is syntenic to *REF6/AtPKDM9A* (Supplemental Figure 2). Thus, we named *MtrunA17Chr3g0116441* and *MtrunA17Chr1g0199511* genes as *MtPKDM9A* and *MtPKDM9B,* respectively. A multiple alignment of the amino acid sequence of MtPKDM9B with the sequences of AtPKDM9B/ELF6 and their counterparts in the legume species *Lotus japonicus*, common bean (*Phaseolus vulgaris*) and soybean (*Glycine max*) revealed a high degree of sequence conservation among these proteins, particularly in the JmjN, JmjC and Cys2His2 zinc finger (zf-C2H2) domains (Supplemental Figure 3).

The TRAP-seq analysis revealed that *MtPKDM9B* gene produces two AS variants, *MtPKDM9B.1* and *MtPKDM9B.2* (Figure 1A). The *MtPKDM9B.1* transcript variant encodes the full-length protein containing a JmjN terminal domain, a JmjC terminal domain and four ZF-C2H2, whereas skipping of exon 3 in the *MtPKDM9B.2* variant introduces a premature stop codon at the end of the JmjC domain (Figure 1B). Quantification of these *MtPKDM9B* transcript variants revealed that none of them are regulated at the level of total RNA abundance at 48 hours post-infection (hpi) (Figure 1C). This is in agreement with previously reported RNA-sequencing data of spot-inoculated roots (Schiessl et al., 2019), in which *MtPKDM9B* transcript levels do not vary between 2 and 120 hpi with *S. meliloti*; however, they are significantly up-regulated at later time points (7 dpi) when nodules are already formed (Supplemental Figure 4). Interestingly, TRAP-seq analysis revealed that *MtPKDM9B* transcript variants are oppositely modulated at translational level in response to rhizobia at 48 hpi, with *MtPKDM9B.1* increasing and *MtPKDM9B.2* decreasing their association to translating ribosomes (Figure 1C). Reverse transcription followed by PCR experiments (RT-PCR) using primers that anneal to exons 2 and 4 verified the occurrence of these two AS transcript variants, as well as their differential regulation at translational level upon rhizobia infection (Figure 1D). Alternative splicing due to alternative 3’acceptor site and 5’donor site was also observed for the two putative orthologs of *MtPKDM9B* in soybean, *Glyma.20G181000* and *Glyma.10G209600*, respectively (Supplemental Figure 5). Analysis of public soybean translatomes (Sainz et al., 2022) revealed that these alternative transcripts are also differentially associated with translation machinery during nodulation (Supplemental Figure 5). suggesting that differential translation of alternative splicing variants, although due to different alternative splicing events, is a shared mechanism present in different legume plants that could contribute to the regulation of expression of this putative lysine demethylase during symbiosis.

**Figure 1.**
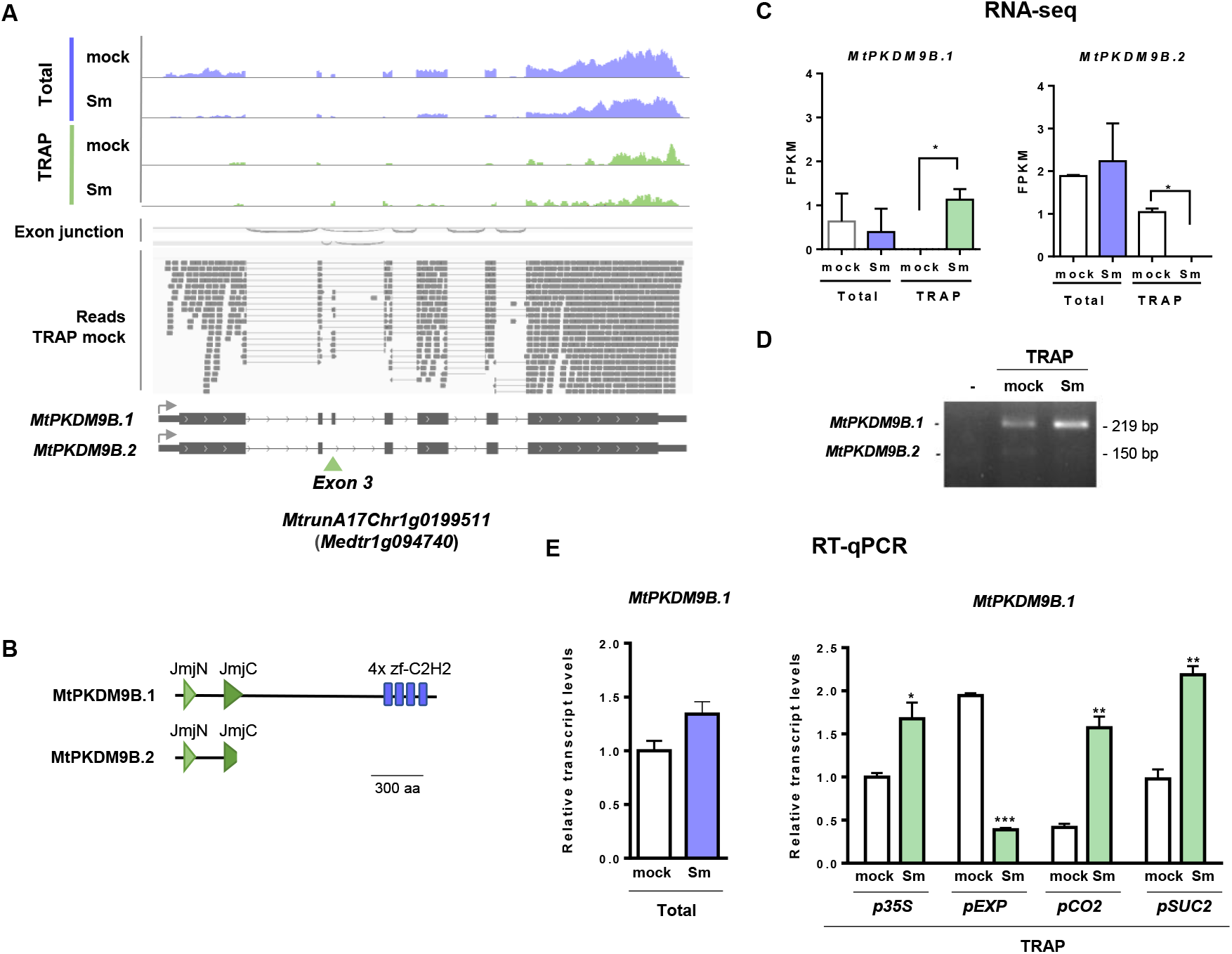
*MtPKDM9B* alternative spliced transcript variants are differentially modulated during early stages of the root nodule symbiosis. **(A)** Distribution of RNA-seq reads of *MtPKDM9B* transcript variants in mock-inoculated or *S. meliloti*-inoculated (Sm) total RNA (blue) or TRAP RNA (green) samples. Exon junctions and reads from TRAP mock samples illustrating skipping or inclusion of exon 3 are presented. The gene model and its transcript variants are schematized below. The gray arrow indicates the transcriptional start site. The green arrowhead indicates the exon skipping event that differentiates *MtPKDM9B.1* and *MtPKDM9B.2* transcripts. **(B)** Scheme of the protein domains present in the products of *MtPKDM9B.1* and *MtPKDM9B.2* transcript variants. The JmjN (light green triangle), JmjC (dark green triangle) and zinc fingers (blue rectangles) domains are schematized. **(C)** RNA-seq quantification of *MtPKDM9B.1* and *MtPKDM9B.2* transcript variants in Total and TRAP RNA samples from mock and Sm inoculated roots. FPKM: fragments per kb per million reads. **(D)** *MtPKDM9B.1* and *MtPKDM9B.2* transcript variants amplified by RT-PCR using primers flanking exon 3 in RNA samples of mock and Sm inoculated roots. Expected sizes of the amplified fragments correspond to 219 bp and 150 bp for *MtPKDM9B.1* and *MtPKDM9B.2,* respectively. **(E)**. RT-qPCR validation of the levels of the *MtPKDM9B.1* variant in Total RNA samples extracted from mock (white bar) and *S. meliloti* roots (blue bar) expressing FLAG-RPL18 under the *p35S* promoter (*p35S*) or in TRAP samples from mock (white bars) and Sm inoculated roots (green bars) expressing the FLAG-RPL18 protein in almost all root cells (*p35S*), epidermal cells (*pEXP7*), cortical cells (*pCO2*), or phloem companion cells (*pSUC2*). Expression values were normalized with *HIS3L* transcript values and expressed relative to the p35S mock sample. Each bar represents the mean ± standard error (SEM) of two biological replicates, with three technical replicates each. Asterisks indicate statistically significant differences in an unpaired two-tailed Student’s t-test (*: pP≤0.05, **: pP≤0.01, ***: pP≤0.001).

TRAP-qPCR was also applied to explore translational regulation in specific tissues (Traubenik et al., 2020). RT-qPCR analysis of TRAP samples from root-specific tissues using primers that specifically detect the *MtPKDM9B.1* variant revealed that this variant increased its association with the translational machinery in developing cortical cells and phloem companion cells but decreased in the epidermal cell layer upon infection with rhizobia (Figure 1E). In agreement, a single nuclei gene expression atlas of *M. truncatula* roots (Cervantes-Perez et al., 2022) indicated that *MtPKDM9B* mRNA levels increased in developing cortical cells and phloem at 48 hpi with *S. meliloti* (Supplemental Figure 6).

### Knockdown of *MtPKDM9B* alters root architecture under nitrogen availability

To explore the function of *MtPKDM9B,* a post-transcriptional gene silencing using an RNA interference (RNAi) approach was applied. The RNAi fragment was designed to target the fourth exon and thus intended to silence both transcript variants of *MtPKDM9B*. In addition, we generated an artificial miRNA targeting the third exon of *MtPKDM9B* (amiR *MtPKDM9B)* to specifically down-regulate the *MtPKDM9B.1* transcript variant (Figure 2A). RT-qPCR with primers that detect both *MtPKDM9B* variants indicated that expression of the *MtPKDM9B* RNAi fragment reduced mRNA levels of both *MtPKDM9B* variants by nearly 50% as compared with roots transformed with a *GUS* RNAi used as a control, but not mRNA levels of its paralog *MtPKDM9A* (Figure 2B). On the other hand, RT-qPCR using primers that specifically detected the *MtPKDM9B.1* variant revealed that expression of the amiR *MtPKDM9B* reduced levels of the *MtPKDM9B.1* by more than 80% as compared with roots transformed with the empty vector (EV) used as a control (Figure 2C). When primers that detected both variants were used, a not significant reduction (lower than 40%) in total *MtPKDM9B* levels were observed, indicating that the knockdown caused by amiR *MtPKDM9B* seems to be specific to the *MtPKDM9B.1* variant (Figure 2C). In addition, expression of the amiR *MtPKDM9B* had no impact on mRNA levels of *MtPKDM9A* (Figure 2C).

**Figure 2.**
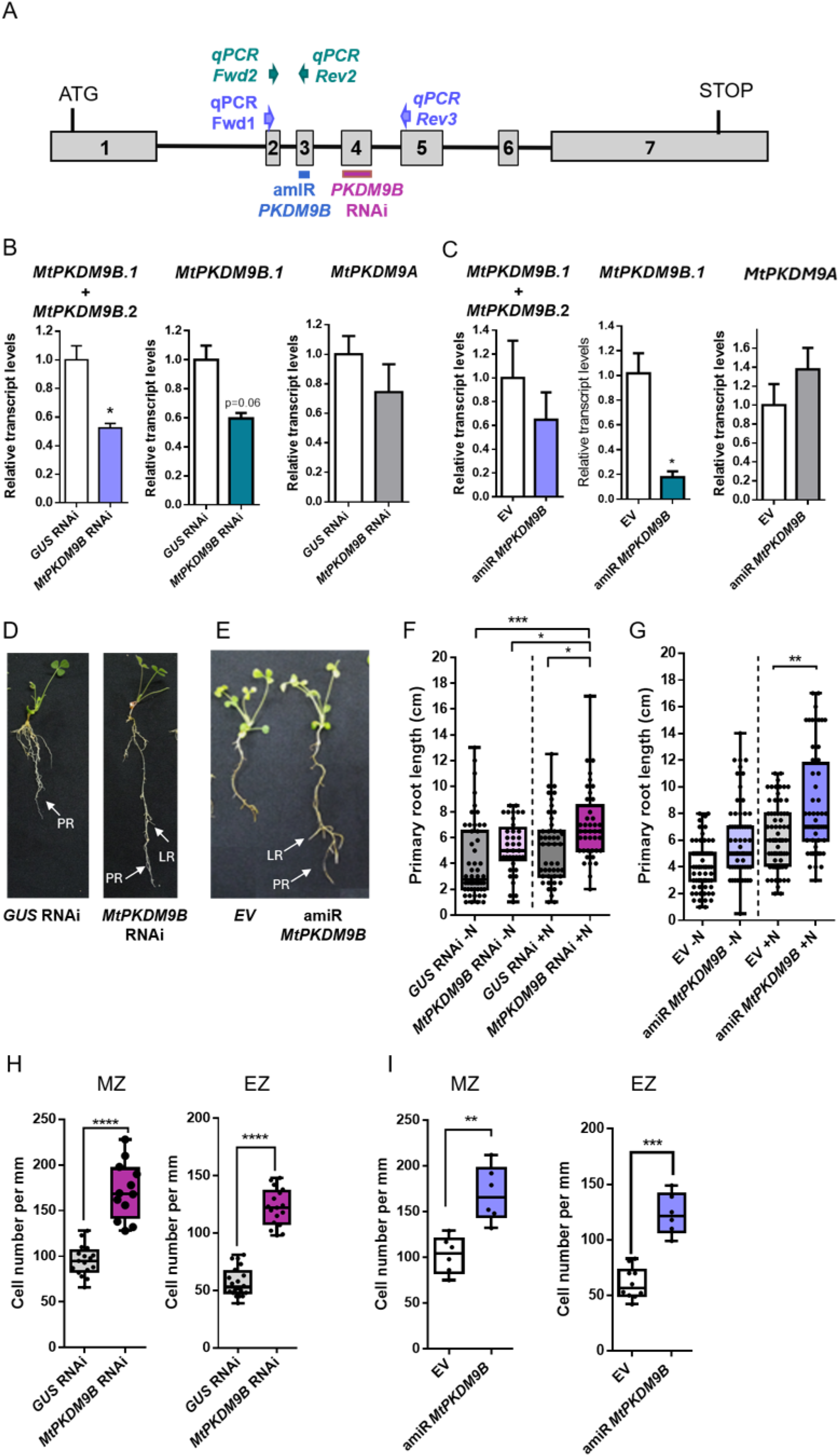
Knockdown of *MtPKDM9B* promotes primary root growth. **(A)** Scheme of the *MtPKDM9B* gene, where introns are represented by lines and exons by light gray rectangles. The position of the RNAi fragment and the amiRNA, as well as primers used in qPCR are indicated. **(B-C)** Relative levels of total *MtPKDM9B* transcripts (*MtPKDM9B.1* and *MtPKDM9B.2* variants), the *MtPKDM9B.1* variant or *MtPKDM9A* in roots of *GUS* RNAi and *MtPKDM9B* RNAi roots (B) or in EV and amiR *MtPKDM9B* roots **(C)**. Transcript levels were quantified by RT-qPCR, normalized by the *HIS3L* transcript values and expressed relative to *GUS* RNAi **(B)** or EV **(C)** samples. Bars are the mean of three biological replicates and error bars represent SD. The asterisk indicates statistically significant differences with p<0.05 (*) in a two-tailed unpaired Student’s t-test. **(D-E)** Representative pictures illustrating root growth of *MtPKDM9B* RNAi **(D)**, and amiR *MtPKDM9B* **(E)** and their controls, *GUS* RNAi and EV, respectively, at 14 days after transformation. **(F-G)** Primary root length of *MtPKDM9B* RNAi roots **(F)**, and amiR *MtPKDM9B* roots (G) and their controls at 14 days after transplantation to media free of nitrogen (-N) or supplemented with KNO_3_ (+N). Boxes extend from the 25th to 75th percentiles, the middle line is the median, and whiskers extend to the minimum and maximum values of three technical replicates with at least 35 plants each. Asterisks denote statistically significant differences in an unpaired two-tailed Student’s t-test (∗∗∗p ≤ 0.001 and ****p ≤ 0.0001). **(H-I)** Cell number per root mm in the meristematic zone (MZ) and elongation zone (EZ) of the roots *GUS* RNAi and *MtPKDM9B* RNAi roots (H) and EV and amiR *MtPKDM9B* roots **(I)**. Asterisks denote statistically significant differences in an unpaired two-tailed Student’s t-test (∗∗p ≤ 0.01, ∗∗∗p ≤ 0.001 and ****p ≤ 0.0001).

During our transformation experiments, we noticed that roots silenced in *MtPKDM9B* by either RNAi or amiR exhibited longer primary roots (Figure 2D and 2E). Thus, we investigated whether the root architecture was modified in roots that express either the *MtPKDM9B* RNAi or the amiR *MtPKDM9B* under sufficient nitrogen supply or under nitrogen starvation. Fourteen days after transplantation, both *MtPKDM9B* RNAi and amiR *MtPKDM9B* roots exhibited significantly longer primary roots than *GUS* RNAi and EV roots, respectively, when plants were grown under sufficient nitrogen supply (Figure 2F and 2G). The longer primary root phenotype was also observed at a later point, i.e., 21 days after transplantation when plants grew on sufficient nitrogen (Supplemental Figure 7). However, no significant differences were observed in roots subjected to nitrogen starvation (Figure 2F and 2G). These results indicate that *MtPKDM9B* might function as a negative regulator of primary root growth, and that this function depends on nitrogen availability. A higher density of lateral roots was also observed in *MtPKDM9B* RNAi roots as compared with *GUS* RNAi roots in plants grown under sufficient or limited nitrogen supply, but not in amiR *MtPKDM9B* as compared with EV roots at 14 days after transplantation (Supplemental Figure 8). On the other hand, the length of lateral roots was not altered by expression of *MtPKDM9B* RNAi and amiR *MtPKDM9B* under sufficient or limited nitrogen supply (Supplemental Figure 8). Since the length of the primary roots increased by the silencing of *MtPKDM9B* in the presence of nitrogen, we measured cell number in the meristematic and elongation zones of roots stained with propidium iodide. The number of cells was higher in both zones of the *MtPKDM9B* RNAi and amiR *MtPKDM9B* roots as compared with those of *GUS* RNAi and EV roots, respectively (Figure 2H and 2I), indicating that *MtPKDM9B* modulates primary root growth by controlling cell division in the root meristem and elongation zone when there is sufficient nitrogen available.

### *MtPKDM9B* mediates infection by rhizobia and nodule development

*MtPKDM9B* transcript variants were differentially regulated at translational level at early stages of the root nodule symbiosis; thus, we explored whether silencing of these variants altered nodule formation or development. The knockdown of both *MtPKDM9B* variants using RNAi did not affect the number of nodules formed over the time as compared with *GUS* RNAi roots (Figure 3A). However, the specific knockdown of the *MtPKDM9B.1* variant using amiR *MtPKDM9B* caused significant reduction of nodule number as compared with EV roots at all times analyzed (7, 10, 14 and 21 dpi) (Figure 3B). These apparently contrasting results might be explained by the lower silencing of the full-length coding transcript variant *MtPKDM9.1* caused by the RNAi construct (Figure 2B), which was insufficient to affect nodule number. In addition, the expression of *MtPKDM9B* RNAi or the amiR *MtPKDM9B* significantly reduced by 35% or 40% nodule size as compared with *GUS* RNAi or EV, respectively (Figures 3C-3F). Moreover, inoculation with a *S. meliloti* strain that constitutively expresses the red fluorescent protein (RFP) revealed that nodule occupancy was reduced by silencing of *MtPKDM9B* (Figures 3G and 3H). Therefore, we evaluated the number of infection events and their progression to the cortical cells. Silencing of *MtPKDM9B* using RNAi or amiR *MtPKDM9B* significantly reduced the density of infection events by nearly 40% in comparison with *GUS* RNAi or EV roots, respectively (Figures 3I and 3J). In addition, the progression of the infection events was slightly delayed in *MtPKDM9B* RNAi roots or amiR *MtPKDM9B*, with a higher percentage of infection events that remained at the stage of microcolony and a lower percentage of the infection events that reached cortical cells (Figures 3K and 3L).

**Figure 3.**
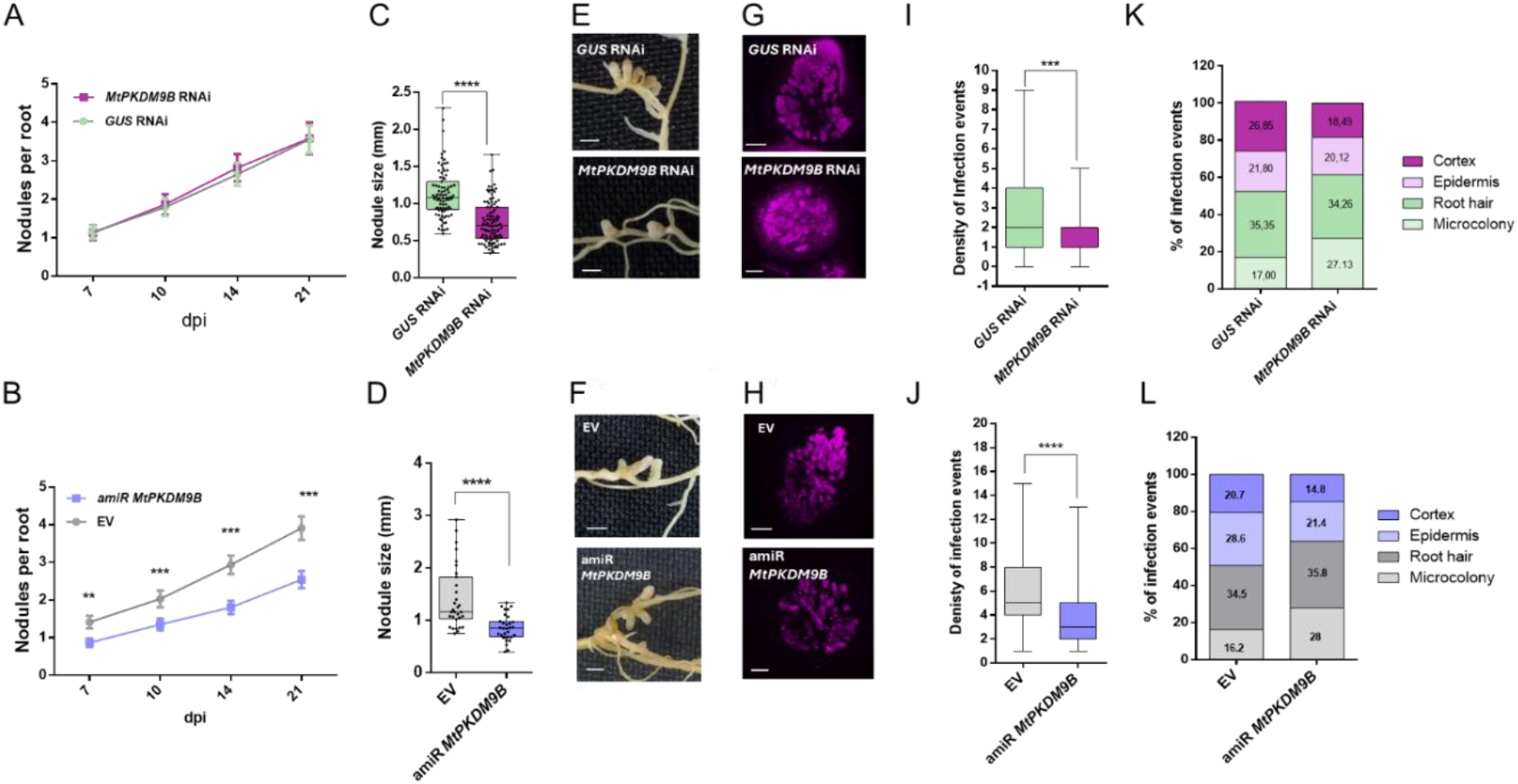
Knockdown of *MtPKDM9B* affects nodule development and infection by rhizobia. **(A-B).** Nodules formed at 7, 10, 14, and 21 dpi with the *S. meliloti* in *MtPKDM9B* RNAi and *GUS* RNAi **(A)** or with amiR *MtPKDM9B* and EV **(B)** roots. Means of three biological replicates are presented for each time point. Error bars represent standard error of the mean (SE). Asterisks indicate statistically significant differences in an unpaired two-tailed Student’s t-test (∗∗p ≤ 0.01 and ∗∗∗p ≤ 0.001). **(C-D)** Nodule size measured at 21 dpi as the length from the base of the nodule to the apex, in *MtPKDM9B* RNAi and *GUS* RNAi **(C)** or in amiR *MtPKDM9B* and EV **(D)**. Bars are the means of three biological replicates and error bars represent the SEM. ∗∗∗ indicates statistically significant differences between samples in an unpaired two-tailed Student’s t-test with p ≤ 0.001. **(E-F)**. Picture of 21 dpi nodules developed in *MtPKDM9B* RNAi and *GUS* RNAi **(E)** or in amiR *MtPKDM9B* and EV **(F)** roots. Bars: 1 mm. **(G-H)** Nodule occupancy by a *S. meliloti* strain expressing RFP (Red fluorescent protein) in *MtPKDM9B* RNAi and *GUS* RNAi **(G)** or in amiR *MtPKDM9B* and EV **(H)** roots. Bars: 50 µm. **(I-J)** Density of infection events (number of infection events per root cm) at 7 dpi with a *S. meliloti* strain expressing the RFP protein in *MtPKDM9B* RNAi (I) or in amiR *MtPKDM9B* roots. **(J)**. Boxes extend from the 25th to 75th percentiles, the middle line is the median and whiskers extend to the minimum and maximum values of three biological replicates, each with more than 30 transgenic roots. Asterisks indicate statistically significant differences in an unpaired two-tailed Student’s t-test (**p ≤ 0.001, ****p ≤ 0.0001). **(K-L)** progression of infection events in *MtPKDM9B* RNAi **(K)** or in amiR *MtPKDM9B* **(L)** roots. Infection events were classified as microcolonies, infection threads (ITs) that end in the root hair, in the epidermal cell layer, or reach the cortex at 7 dpi. Each category is presented as the percentage of total infection events. Data are representative of three independent biological replicates, each with more than 25 transgenic roots.

The reduction of the nodule size and infection events suggest that the nodules formed in *MtPKDM9B* silenced roots might not be functional in nitrogen fixation. Thus, we evaluated the viability of the bacteria within the nodules by a live/death assay that uses the green fluorescent dye SYTO9 to stain live bacteria accommodated in nodule cells and the red fluorescent dye propidium iodide to stain cells with membrane damage within the infection and fixation zones of the nodule, as well as meristematic cells in the nodule apex. Confocal microscopy indicated that silencing of both *MtPKDM9B* variants or the *MtPKDM9B.1* variant using either RNAi or amiR, respectively, significantly reduced the viability of the bacteria within the nodule since most of the cells formed in *MtPKDM9B* RNAi and amiR *MtPKDM9B* nodules were stained with propidium iodide, whereas the majority of the cells in *GUS* RNAi and EV nodules were stained with SYTO9 (Figure 4A-D). Consistently, transcript levels of the leghemoglobin *MtLGHB1* gene were reduced by nearly 80% in nodules formed in *MtPKDM9B* RNAi or amiR *MtPKDM9B* roots as compared with those formed in *GUS* RNAi or EV roots, respectively (Figure 4C and 4D).

**Figure 4.**
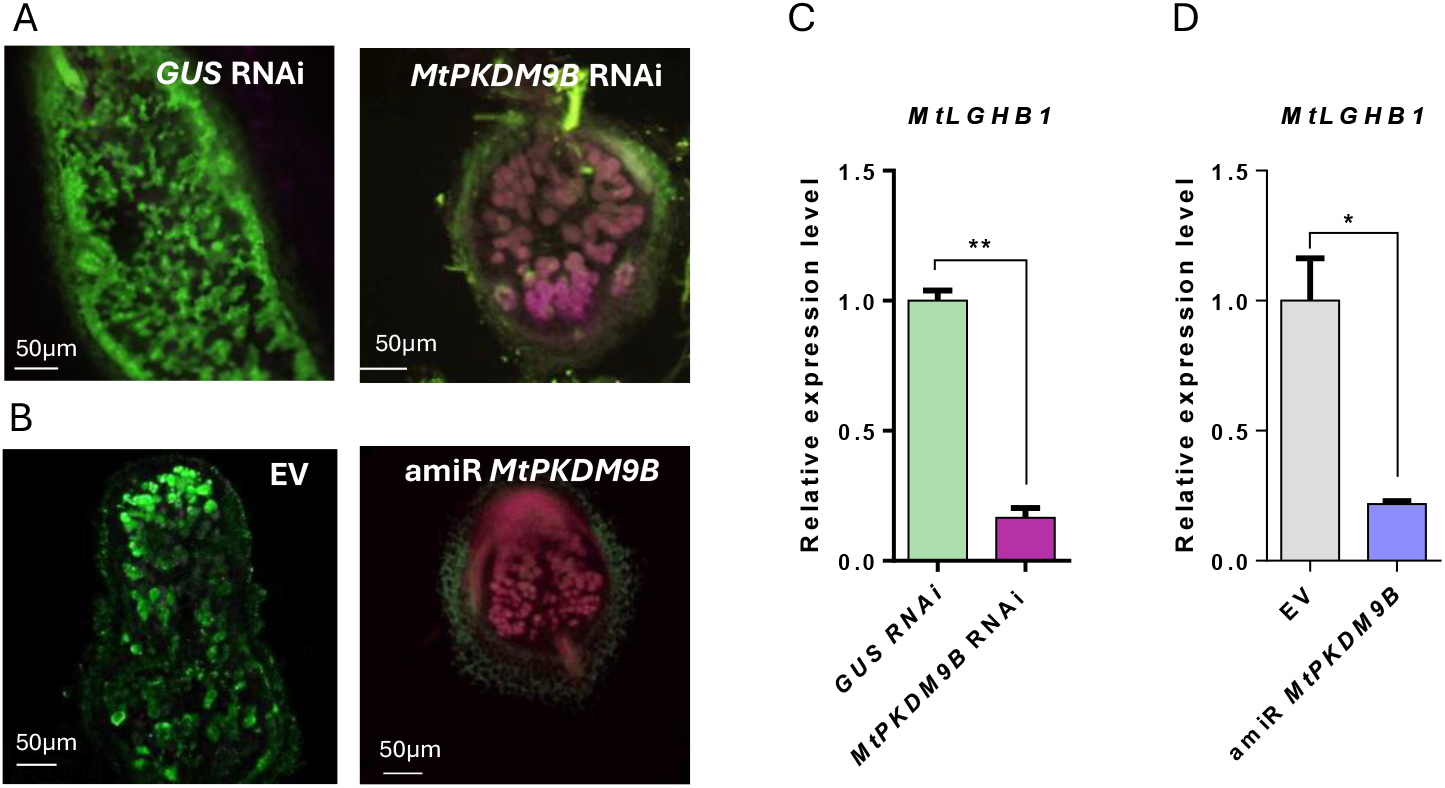
Knockdown of *MtPKDM9B* affects bacterial viability within nodules and expression of the leghemoglobin gene *MtLGHB1.* **(A-B)** Live/dead staining of nodules at 21 dpi with SYTO9 (green), which stains live bacteria, and propidium iodide (magenta), which is incorporated into cells with damaged plasma membranes and into the nuclei of meristematic cells in *GUS* RNAi and *MtPKDM9B* RNAi (A) or EV and amiR *MtPKDM9B* **(B)** roots. Bar = 50 μm. **(C-D)** Relative expression of *MtLGHB1* in nodules of 21 dpi of *GUS* RNAi and *MtPKDM9B* RNAi **(C)** or EV and amiR *MtPKDM9B* **(D)** roots. *MtLGHB1* transcript levels were quantified by RT-qPCR. Values were normalized by the *HIS3L* transcript values and expressed relative to *GUS* RNAi **(C)** or EV **(D)** samples. Bars are the mean of three biological replicates and error bars represent standard deviation (SD). The asterisks indicate statistically significant differences in a two-tailed unpaired Student’s t-test (∗p ≤ 0.05 and ∗∗p ≤ 0.01).

### Knockdown of *MtPKDM9B* alters H3K27me3 landscape in *M. truncatula* roots during symbiosis

ELF6, the putative ortholog of MtKDM9B in Arabidopsis, has histone demethylase activity on H3K27me3; moreover, *elf-6* mutants exhibited altered H3K27me3 homeostasis (Antunez-Sanchez et al., 2020). Thus, we questioned whether silencing of *MtPKDM9B* could alter H3K27me3 landscape in *M. truncatula* roots. Western blot analysis with an α-H3K27me3 antibody revealed a significant increase in H3K27me3 levels in *MtPKDM9B* RNAi roots as compared to *GUS* RNAi in both mock and *S. meliloti* inoculated conditions (Figure 5A). To investigate the effect of *MtPKDM9B* silencing on the distribution of H3K27me3 marks at genome wide scale, chromatin immunoprecipitation followed by RNA sequencing (ChIP-seq) assays were performed in *GUS* RNAi and *MtPKDM9B* RNAi roots under mock and *S. meliloti* inoculated conditions using a commercial α-H3K27me3 antibody or α-rabbit IgG as a control (Supplemental Figure 9A). Induction of the symbiotic marker *Ethylene Response Factor Involved in Nodulation 1* (*MtERN1*) in response to rhizobia was verified prior to the ChIP-seq experiment in both *MtPKDM9B* RNAi and *GUS* RNAi roots (Supplemental Figure 9B). To evaluate whether ChIP samples were enriched in regions with high H3K27me3 levels, qPCR were performed on ChIP samples using primers designed to detect a locus containing a repetitive region (*MtrunA17Chr8R0004680*) described as marked with H3K27me3 in root tissues in a previous report (Pécrix et al., 2018). The results verified the enrichment of the H3K27me3 mark in this locus on ChIP samples from both *GUS* RNAi and *MtPKDM9B* RNAi roots (Supplemental Figure 9C). Differential analysis was performed on ChIP-seq data to identify differential H3K27me3 regions (DHMRs) in the different pairwise comparisons (|Log_2_Fold Change (FC)|≥ 1, p-value≤0.05). Comparison of *S. meliloti* and mock *GUS* RNAi samples identified 1126 hypermethylated and 6340 hypomethylated DHMRs representing 2.7% of the *M. truncatula* genome (Supplemental Table 1). These DHMRs were mainly located in gene bodies (32%), intergenic (50%), promoter (14%) and downstream regions (4%) (Figure 5B and 5C). Many of these DHMRs are located within or near symbiotic genes known to be upregulated at 48 hpi in response to *S. meliloti* (Schiessl et al., 2019), including*, MtNOOT1* (*Nodule root 1*) (Couzigou et al., 2012) and *MtNIN-like protein* (*MtNLP1*) (Luo et al., 2023), as well as numerous genes encoding NCR (Nodule Cysteine Rich) peptides (Supplemental Table 1), some of which are required for bacteria differentiation (Van de Velde et al., 2010). The comparison between *S. meliloti* inoculated and mock *MtPKDM9B* RNAi roots identified 21185 hypermethylated and 24283 hypomethylated DHMRs representing 20.7% of the *M. truncatula* genome (Supplemental Table 2), which were distributed in gene bodies (38%), intergenic (44%), promoter (14%) and downstream (4%) regions (Figure 5B and 5C,). The number of DHMRs was much higher in *MtPKDM9B* RNAi roots than in *GUS* RNAi roots, highlighting the global deregulation of changes in H3K27me3 levels in response to rhizobia caused by silencing of *MtPKDM9B* (Figure 5B). A very low percentage (0.7%, 159 DHMRs) of the hypermethylated DHMRs in response to *S. meliloti* inoculation in *MtPKDM9B* RNAi roots were also hypermethylated in *GUS* RNAi roots, whereas 5.3% (1293 DHMRs) of hypomethylated regions in *MtPKDM9B* RNAi were also hypomethylated in *GUS* RNAi in response to *S. meliloti* inoculation (Figure 5D), reinforcing the notion that silencing of *MtPKDM9B* has profound effects on the H3K27me3 landscape at early stages of the root nodule symbiosis. Among hypomethylated DHMRs in response to *S. meliloti* in both *GUS* RNAi and *MtPKDM9B* RNAi roots were DHMRs contained within genes involved in the control of nodulation such as *MtNLP1* and *MtNOOT1* (Figure 5E).

**Figure 5.**
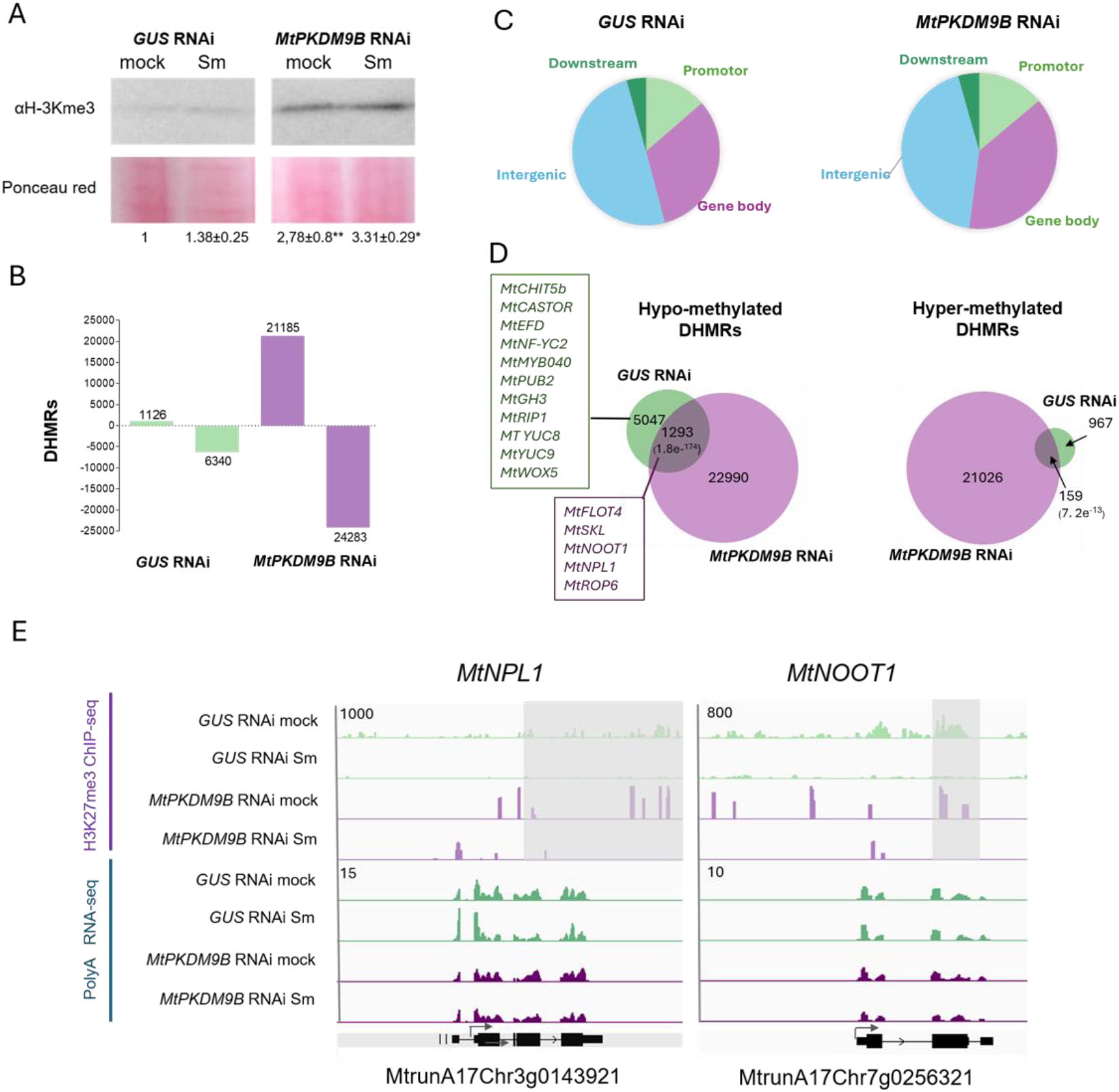
Silencing of *MtPKDM9B* alters H3K27me3 in mock and *S. meliloti* inoculated roots. **(A)** Western blot of *GUS* RNAi and *MtPKDM9B* RNAi nuclear extracts of mock and *S. meliloti* (Sm) inoculated roots using α-H3K27me3 antibodies. Lower panel presents the Ponceau red staining. Numbers indicate the quantification of three independent Western blots. **(B)** Differentially H3K27me3 regions (DHMRs) identified by ChIP-seq on *GUS* RNAi and *MtPKDM9B* RNAi roots at 48 hpi with Sm using the *diffreps* tool with a 1000-bp window, a 1<Log_2_FC<-1 and a p-value< 0.05. Positive and negative bars represent hyper- and hypo-methylated DHMRs, respectively. **(C)** Pie charts showing the distribution of DHMRs in gene bodies, promoter, intergenic and downstream regions identified using ChIPeakAnno tool **(D)** Venn diagrams showing the overlap between hypo- and hyper-methylated DHMRs in *GUS* RNAi and *MtPKDM9B* RNAi roots at 48 hpi with Sm. **The p-value for Fisher’s exact test is shown in brackets (E) Integrative genomic viewer (IGV) captures of** ChIP-seq signals with H3K27me3 and RNA-seq normalized reads for the *MtNPL1* and *MtNOOT1* genes in mock and Sm inoculated *GUS* RNAi (green) and *MtPKDM9B* RNAi (purple) root samples. Normalized reads are presented. Shade gray boxes are DHMRs that are hypomethylated in Sm samples as compared to mock samples for each gene. Gene models are presented below, and arrows indicate the transcriptional start site (TSS). Numbers on the top left indicate the maximum read value of the scale, which was the same for each gene in all ChIP-seq or RNA-seq samples.

The comparison of DHMRs hypomethylated in response to rhizobia in *GUS* RNAi, but not in *MtPKDM9B* RNAi roots yielded 5,047 DHMRs contained within or near 4,035 genes (Figure 5D, Supplemental Table 3). These DHMRs were considered putative targets of MtPKDM9B during symbiosis since demethylation of these loci at 48 hpi with rhizobia requires *MtPKDM9B*. Comparison with genes located in symbiotic island (Pécrix et al, 2018) revealed that 4.5% of the MtPKDM9B targets were located in symbiotic islands. Gene ontology analysis revealed that the most prominent biological process categories were “DNA binding”, “Regulation of transcription” and “Transcription factor activity” (Supplemental Figure 10A). The list includes key genes involved in early nodulation signaling and infection such as *MtCASTOR* (Ane et al., 2004), *MtMYB040* (Gao et al., 2026) and *MtRIP1* (*Rhizobia Induced Peroxidase 1*) (Cook et al., 1995) (Supplemental Figure 10B), as well as genes functioning in nodulation that have been recruited from root developmental programs such as *MtPLT4/MtBBM* (Franssen et al., 2015)*, MtWOX4 and MtWOX5* (Osipova et al., 2012) (Figure 5D, Supplemental Figure 10C, Supplemental Table 3).

We also compared DHMRs in *MtPKDM9B* RNAi roots versus *GUS* RNAi roots under mock and symbiotic conditions. Under mock conditions, 12,342 hypermethylated DHMRs contained near or within 10,994 genes and 16,073 hypomethylated DHMRs located in 11,054 genes were identified (Supplemental Figure 11A, Supplemental Table 4). Genes with hypermethylated DHMRs in *MtPKDM9B* RNAi roots as compared with *GUS* RNAi included members of the ARF family including *MtARF2*, *MtARF3*, *MtARF4* (Kirolinko et al., 2021) and *MtARF16* (Breakspear et al., 2014), which were involved in the control of root architecture and nodule development, genes encoding cyclins from the CycA, CycB and CycD families, four NLP genes, which play roles in nitrate-induced inhibition of nodulation (Luo et al., 2021) and more than 10 members of the NPF (Nitrate and Peptide Transporter Family) (Morère-Le Paven et al., 2024), which could be related with the root architecture phenotype observed in *MtPKDM9B* RNAi roots in the presence of nitrogen (Figure 2D-2F). Under symbiotic conditions, we identified 15,845 hypermethylated DHMRs located near or within 11939 genes and 8572 hypomethylated DHMRs located near o within 6781 genes (Supplemental Figure 11A, Supplemental Table 5). Genes with hypermethylated DHMRs showed significant enrichments in the GO categories of “DNA binding”, “RNA Pol II transcription factor activity-sequence specific DNA binding”, “Transcriptional regulation”, “Post-transcriptional regulation”, “Protein binding and protein-protein interactions”, as well as “Hormone-mediated signaling pathway” and “Nucleosome dependent ATPase activity” (Supplemental Figure 11B). Notably, *MtPKDM9B* RNAi versus *GUS* RNAi roots under symbiotic conditions showed hypermethylated DHMRs in key genes for early signaling and rhizobia infection including *MtERN2* (Andriankaja et al., 2007; Cerri et al., 2012; Cerri et al., 2016)*, MtCASTOR* (Ane et al., 2004), *MtCNGC15a* and *MtCNGC15c* (*Cyclic Nucleotide-Gated Channel 15a and 15b*) (Charpentier et al., 2016), *MtNENA* (Groth et al., 2010)*, MtNSP1* (Smit et al., 2005) and *MtCYCLOPS/MtIPD3 (Interacting protein with DMI3)* (Limpens et al., 2011); genes required for nodule organogenesis and development such as *MtEFD2* (Jardinaud et al., 2022), *MtCCS52b* (Tarayre et al., 2004), *MtNF-YB18* (Baudin et al., 2015) and *MtSCR* (Dong et al., 2021), but also genes involved in auxin signaling and auxin biosynthesis, which have been recruited from root developmental programs to act in the root nodule symbiosis, including *MtLBD16* (*Lateral Organ Boundaries Domain 16*), *MtSTYL1* (*SHORT INTERNODES/STYLISH 1*), and *MtYUC9* (Schiessl et al., 2019; Soyano et al., 2019) (Supplemental Table 5). To validate differential H3K27me3 methylation detected by ChIP-seq we selected three genes that are highly methylated in rhizobia inoculated *MtPKDM9B* RNAi but not in *GUS* RNAi roots, i.e., *MtLBD16*, *MtNSP1* and *MtSTYL1* (Supplemental Figure 12A), and conducted ChIP-qPCR assays using primers flanking the DHMR identified for each gene. The results verified that H3K27me3 levels in these loci remained high or increased upon rhizobia inoculation in *MtPKDM9B* RNAi roots (Supplemental Figure 12B).

### Knockdown of *MtPKDM9B* affects the expression of genes required for infection by rhizobia and nodule development

To further understand the function of *MtPKDM9B* and the relevance of MtPKDM9B-mediated demethylation of H3K27me3 for regulation of gene expression during symbiosis, we performed RNA-seq experiments in *GUS* RNAi and *MtPKDM9B* RNAi roots under mock and *S. meliloti* inoculated conditions. Principal Component Analysis (PCA) and correlation analysis revealed that *MtPKDM9B* RNAi samples from *S. meliloti* inoculated roots were closer to mock inoculated *GUS* RNAi and *MtPKDM9B* RNAi samples than to *S. meliloti* inoculated *GUS* RNAi samples (Supplemental Figure 13), indicating that knockdown of *MtPKDM9B* impairs the transcriptional response to rhizobia. This is also supported by the smaller number of differentially expressed genes (DEGs) up or down regulated in *MtPKDM9B* RNAi roots (188 or 27 DEGs, respectively) as compared to *GUS* RNAi roots (1969 or 1549, respectively) using a |Log_2_FC|>1, Padj < 0.05 criteria (Figure 6A, Supplemental Table 6). Common DEGs up regulated in response to rhizobia in both *GUS* RNAi and *MtPKDM9B* RNAi roots include several symbiotic markers, e.g., *MtENOD11, MtENOD12, MtERN1* and *MtNIN* among others, whereas other symbiotic genes were significantly up regulated only in *GUS* RNAi roots, including genes mediating rhizobial infection and/or nodule development such as *MtERN3, MtFLOT4, MtLYK3, MtPUB2, and MtGH3* (Figure 6A and 6B). Combination of RNA-seq data with H3K27me3 ChIP-seq data indicated a statistically significant correlation (Fisher’s exact test) between H3K27me3 hypermethylated genes and DEGs down regulated in *MtPKDM9B* RNAi roots as compared with *GUS* RNAi roots under both mock and *S. meliloti* inoculated conditions, whereas no significant correlation was observed between hypomethylated genes and up regulated DEGs in the same comparison (Supplemental Figure 14). Interestingly, a significant proportion (136 genes, representing 7.5%) of the DEGs up regulated in response to rhizobia only in *GUS* RNAi roots were also hypomethylated in response to rhizobia in *GUS* RNAi roots but not in *MtPKDM9B* RNAi roots, including *MtPUB2, MtCHIt5b,* and *MtGH3,* among others (Figure 6C and 6D). In addition, we identified the recently described gene *MtMYB040* (Gao et al, 2026) as hypomethylated only in *GUS* RNAi roots, whose upregulation in response to rhizobia was lower in *MtPKDM9B* RNAi roots (log_2_FC=1.88, Padj = 6.08 E^−07^) as compared with *GUS* RNAi roots (log_2_FC= 4.26, Padj = 6.32E^−23^) (Figure 6D, Supplemental Table 6). *MtMYB040* encodes a MYB transcription factor interacting with NSP2 to mediate activation of flavonoid biosynthetic genes and control rhizobial infection and nodulation (Gao et al, 2026). These results indicate that knockdown of *MtPKDM9B* has profound impact on H3K27me3 demethylation upon inoculation with rhizobia, which significantly influences the expression of symbiotic genes. Changes in gene expression could be a direct consequence of H3K27me3 levels as in the case of *MtPUB2, MtMYB040, MtCHIT5b* and *MtGH3*, in which knockdown of *MtPKDM9B* impairs demethylation of H3K27me3 and prevents upregulation of gene expression, or indirectly as in the case of *MtERN3*, *MtFLOT4* and *MtLYK3*, in which changes in gene expression were observed without significant changes in H3K27me3 levels as consequence of knockdown of *MtPKDM9B*.

**Figure 6.**
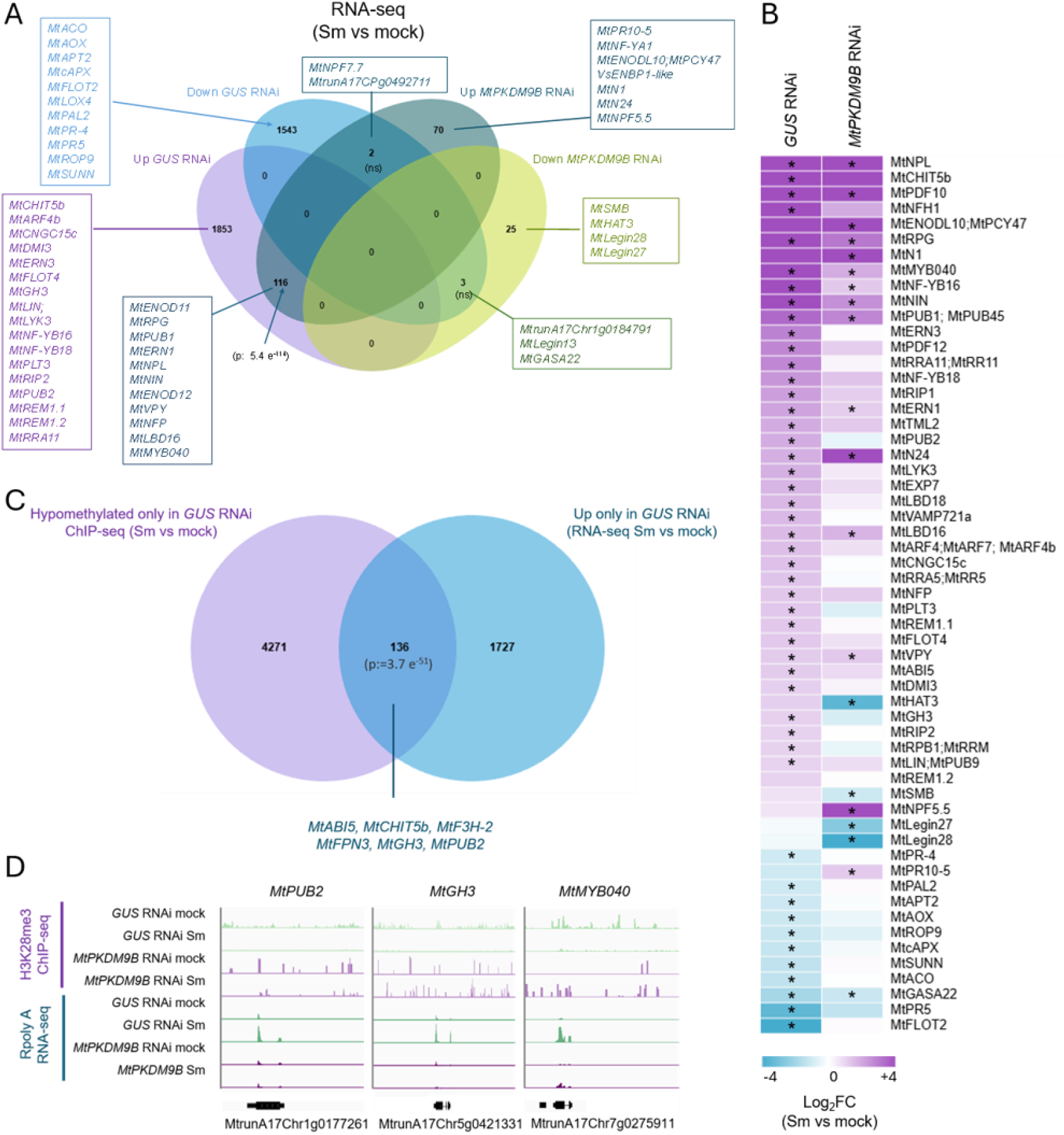
Silencing of *MtPKDM9B* affects the expression of genes required for bacterial infection and nodule development. (**A**) Venn diagrams of DEGs up and down regulated in response *S. meliloti* inoculated root versus mock inoculated roots. Selected DEGs are indicated. P-values for Fisher’s exact test are shown in brackets, ns: not significant. (**B**) Heatmap of selected DEGs. Gene expression levels are depicted as Log_2_FoldChange (FC) in S*. meliloti* (Sm) inoculated versus mock in *GUS* RNAi and *MtPKDM9B* RNAi roots. The asterisk indicates statistically significant differential expression using DEseq2 (Padj < 0.05). (**C**) Venn diagrams showing overlap between genes with hypomethylated H3K27me3 levels and upregulated DEGs in response to *S. meliloti* only in *GUS* RNAi roots. p-value for Fisher’s exact test is shown in brackets. (**D**) Integrative genomic viewer (IGV) captures of subtracted H3K27me3 ChIP-seq signals and RNA-seq signals for *MtPUB2, MtGH3* and *MtMYB040* in *S. meliloti* inoculated (Sm) and mock inoculated *GUS* RNAi and *MtPKDM9B* RNAi roots.

## Discussion

Here, we identified *MtPKDM9B* as a gene encoding a H3K27me3 demethylase with key functions in root development and the establishment of the root nodule symbiosis in *M. truncatula*. *MtPKDM9B* produces two AS transcript variants due to exon skipping, which are oppositely modulated at translational level in response to rhizobia. Interestingly, one of the putative orthologs of *MtPKDM9B* in soybean also produced two alternative transcript variants that are translationally regulated during nodulation (Sainz et al., 2022), suggesting that differential translation of alternative spliced variants in response to rhizobia could be an evolutionary conserved mechanisms in legumes. In animals, it was previously reported that alternative splicing of the histone demethylases encoding gene *LSD1/KDM1* modulates the morphogenesis of neurites in the mammalian nervous system (Zibetti et al., 2010). Additionally, two recent reports showed that alternative splicing of the *UTX/KDM6A* mediates control of mammalian development and diseases (Fotouhi et al., 2023) and that the histone demethylase gene *Kdm6bb* is also subjected to alternative splicing in response to high temperature to mediate sex determination in tilapia (Yao et al., 2023). These studies suggest that alternative splicing of members of the histone demethylase family might be a regulatory mechanism evolutionary conserved in plant and animal kingdoms to mediate development and differentiation.

Phylogenetic and syntenic analysis indicated that the ortholog of *MtPKDM9B* in Arabidopsis is *ELF6* (Noh et al., 2004; Antunez-Sanchez et al., 2020). *ELF6* and its closest paralog *REF6* encode proteins that erase the repressive mark H3K27me3 deposited by the PRC2 in specific tissues, activating the expression of genes involved in flowering (Yan et al., 2018), developmental patterns and the responses to environmental stimuli (Lu et al., 2011). This led us to hypothesize that *MtPKDM9B* could also function in the demethylation of H3K27me3 in *M. truncatula* roots. This was verified through a Western blot analysis revealing that roots silenced in *MtPKDM9B* exhibited greater accumulation of the H3K27me3 mark. Subsequent ChIP-seq data showed that silencing of *MtPKDM9B* altered the landscape of the repressive mark H3K27me3 at specific genomic loci, supporting the notion that the product of *MtPKDM9B* is involved in the removal of repressive H3K27me3 marks, as previously described for its ortholog *ELF6* (Antunez-Sanchez et al., 2020).

Forty-six histone demethylases have been identified in the *M. truncatula* and their expression analyzed at various developmental stages and differentiation zones of nitrogen-fixing nodules (Lopez et al., 2022). However, prior to this study none of them had been functionally characterized during the root nodule symbiosis. Our findings revealed that silencing of the *MtPKDM9B* negatively affects the formation and progression of ITs, indicating that the activity of this H3K27me3 demethylase is required for successful rhizobia infection and subsequent nodule colonization.

Among the *MtPKDM9B* targets that change during symbiosis, i.e. those that decrease their H3K27me3 status in response to inoculation with rhizobia in *GUS* RNAi roots but not in *MtPKDM9B* RNAi roots, we found several genes required for rhizobia infection including *MtCASTOR*, an ion channel that is part of the early signaling of the symbiosis that leads to the calcium oscillations required for the formation of the infection threads (Ane et al., 2004; Peiter et al., 2007), *MtRIP*, a peroxidase that precedes bacterial infection (Cook et al., 1995), *MtPUB2*, a putative ubiquitin ligase that interacts with the LRR-kinase receptor MtDMI2 to mediate rhizobial infection and nodule formation (Liu et al., 2018), *MtGH3*, the putative ortholog of GmGH3, which has been shown to be required for nodule zonation in soybean (Tu et al., 2024) and *MtMYB040*, a gene encoding a MYB transcription factor that inteacts with MtNSP2 to actiavte the expression of genes involved in flavonoid biosynthesis, and is requiered for bacetrial infection and normal levels of nodulation (Gao et al, 2026). In *MtPKDM9B* silenced roots, the loci containing the *MtPUB2*, *MtMYB040* and *MtGH3* genes maintain high levels of the repressive mark H3K27me3 in the presence of rhizobia, impairing their transcriptional activation in response to rhizobia. It was also observed that bacteria within the nodules formed in *MtPKDM9B* silenced plants are not viable at 21 dpi. That is, the few bacteria released into the cortical cells do not reach the differentiation stage that allows them to fix atmospheric nitrogen. This is also reflected in the drastic decrease in the expression of the *MtLGHB1* gene in *MtPKDM9B* silenced roots as compared to control roots. Therefore, *MtPKDM9B* would act not only as a positive regulator of infection, but also as an important modulator ultimately affecting bacteria viability within the nodules. Consistently, ChIP-seq data also identified loci containing genes involved in the control of nodule plant cells and/or bacteroid differentiation that exhibited hypermethylated DHMRs in *MtPKDM9B* RNAi roots as compared to *GUS* RNAi roots under symbiotic conditions. The list includes *MtEFD,* which encodes a transcription factor of the AP2 family that has been involved in the endoreduplication process of plant and bacterial cells, thus affecting the correct differentiation of nodule and bacterial cells (Jardinaud et al., 2022), as well as 45 genes encoding NCR (Nodule-specific Cysteine-Rich) peptides, which participate in differentiation of bacteria into bacteroids, an irreversible transition where bacteria elongate and increase their DNA content through endoreduplication (Mergaert et al., 2006). Most NCRs are expressed in infected nodule tissue and are activated sequentially as nodule organogenesis and bacterial differentiation progress (Guefrachi et al., 2014). Defects in the transcriptional activation of these genes, including the maintenance of the repressive mark H3K27me3 in the presence of rhizobia, could lead to an arrest in bacteroid differentiation and the inability to fix atmospheric nitrogen.

Roots specifically silenced in the long variant *MtPKDM9B.1*, which encodes the full length MtPKDM9B protein, presented a lower number of nodules, whereas silencing of only *MtPKDM9B.1* or both variants affected nodule size, generating less developed nodules than those of their controls. Therefore, it could be possible to speculate that the protein encoded by *MtPKDM9B.1* variant acts as a positive modulator of nodule initiation. These different phenotypes could be explained by the higher association to the translational machinery of the *MtPKDM9B.1* variant in rhizobia inoculated roots, suggesting that this variant could be translated at a higher rate. When *MtPKDM9B.1* was silenced, we observed a more penetrant phenotype than when both variants are silenced. In *MtPKDM9B* RNAi roots only 40% silencing of *MtPKDM9B.1* occurs, and assuming that the observed phenotype is only due to the *MtPKDM9B.1* variant, the less pronounced phenotype could be a consequence of the lower percentage of silencing of this variant.

ChIP-seq data revealed thousands of DHMRs in both *GUS* RNAi and *MtPKDM9B* RNAi roots, many of which are contained near or within symbiotic genes. This is consistent with the higher H3K27me3 levels found in symbiotic island that contain genes required for nodule formation and differentiation (Pecrix et al., 2018). Within this group of genes, we found genes that are essential for nodule organogenesis and development, including *MtNSP1* (Smit et al., 2005), *MtLBD16* and *MtSTYL1* (Schiessl et al., 2019), for which differential methylation was confirmed by ChIP-PCR analysis in independent biological replicates. Some symbiotic genes including *MtPUB2, MtMYB040* and *MtGH3* were hypomethylated in response to rhizobia exclusively in the *GUS* RNAi roots. Moreover, RNA-seq analysis verified up regulation of *MtPUB2* and *MtGH3* and mRNAs in response to rhizobia in *GUS* RNAi root, but not in *MtPKDM9B* RNAi, whereas for MtMYB040 its induction in response to rhizobia was reduced in *MtPKDM9B* RNAi as compared with *GUS* RNAi roots. *MtPUB2* has been shown to be required for rhizobial infection and nodulation (Liu et al., 2018). *MtMYB040* encodes a transcription factor of the MYB family required for bacterial infection and nodulation (Gao et al., 2026), whereas *MtGH3* could be required for nodule differentiation (Tu et al., 2024). Thus, it is possible to speculate that miss regulation of these genes in *MtPKDM9B* RNAi roots could contribute to explaining the phenotype of reduced infection and less developed nodules observed in *MtPKDM9B* silenced roots. Therefore, *MtPKDM9B* could be involved in the removal of the repressive mark H3K27me3 in genomic regions nearby or within these genes that are essential for rhizobial infection and nodule development, activating their transcription at early stages of the root nodule symbiosis (Figure 7). Taking together, the results presented here indicate that the *MtPKDM9B* is subject to translational regulation in cells engaged in the root nodule symbiosis to act as a positive modulator of the symbiosis, mediating demethylation and transcriptional activation of symbiotic genes required for rhizobial infection and the development of functional nitrogen fixing nodules

**Figure 7.**
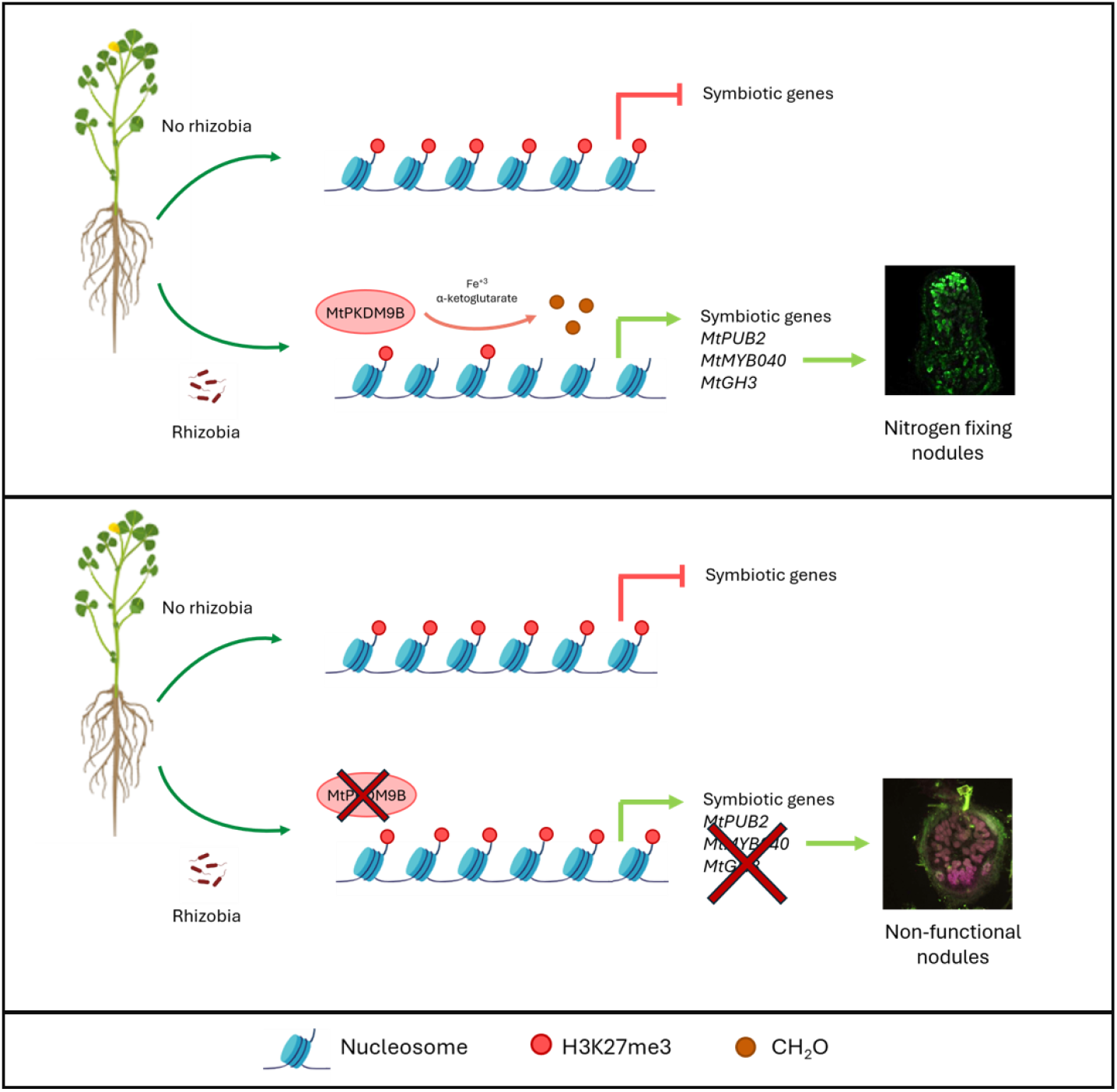
Model proposed for the function of MtPKDM9B in the development of nitrogen fixing nodules. In roots expressing *MtPKDM9B* inoculation with rhizobia enhances translation of *MtPKDM9B.1*, which encodes the full length MtPKDM9B protein. MtPKDM9B-dependent demethylation of H3K27me3 of loci containing symbiotic genes, including *MtPUB2, MtMYB040* and *MtGH3* among others, activates their transcription leading to the formation of functional nitrogen fixing nodules. In roots silenced in *MtPKDM9B*, demethylation of these loci is impaired, preventing transcription of symbiotic genes crucial for bacterial infection and nodule development resulting in non-functional nodules.

In agreement with the fundamental role described for histone methylation in regulating transcriptional reprogramming during plant development and cell differentiation (Morgan and Shilatifard, 2020; Millan-Zambrano et al., 2022), our study revealed that *MtPKDM9B* seems to play a role in root development, since roots silenced in this transcript exhibited a longer root phenotype. REF6 demethylates PCR2-methylated histones at the *CUP-SHAPED COTYLEDON 1* (*CUC1*) gene locus, which is involved in organ boundary formation during seed germination and cotyledon separation (Cui et al., 2016). In root development, the action of the H3K36me3 and H3K9me3 demethylase AtJMJ30 promotes the expression of two members of the LBD transcription factor family, *LBD16* and *LBD19*, stimulating callus formation by establishing root primordium identity (Lee et al., 2018). PRC2 is composed of several proteins that deposit methyl groups on H3K27. Some of these members of the PRC2 exhibit a complex spatial and temporal expression pattern and function in the specification and cell proliferation of the meristem and root vasculature. *clf-28 swn-7* double mutant plants exhibit shorter primary roots with small meristems and increased cell counts in the vascular cylinder as well as a complete loss of H3K27me3 marks (de Lucas et al., 2016). The observation that plants with reduced levels of *MtPKDM9B* exhibited longer PRs suggest that *MtPKDM9B* would act as a negative regulator of primary root growth. This is the opposite effect to that reported for the PRC2 methylases CFL or SWN, which seems to act positively on the growth of primary roots in Arabidopsis (de Lucas et al., 2016). This opposing effect was expected since H3K27me3 methylations produced by PRC2 are antagonized by demethylations caused by ELF6 and REF6 in Arabidopsis (Noh et al., 2004; Antunez-Sanchez et al., 2020). Our data regarding the number of cells at the root apex indicated that the higher root length might be consequence of the increase in cell numbers in the meristematic and elongation zones of the *M. truncatula* root. Increased cell proliferation in *MtPKDM9B* silenced roots could be due to an alteration in the cell cycle control. Our ChIP-seq data indicated that H3K27me3 levels were higher in *MtPKDM9B* silenced roots as compared to *GUS* RNAi roots in regions close to or within genes involved in the cell cycle progression. Among them were those encoding the MtCycA2-3, MtCycBL-3, MtCycC1-1 and MtCycD4-1 (Supplemental Table 4), for which putative orthologs in Arabidopsis have been previously involved in root development during seed germination (Masubelele et al., 2005), and the Cell Cycle Switch 52b (MtCCS52b), which is part of the anaphase activating complex during the G2 to M transition of the cell cycle (Tarayre et al., 2004). In agreement, it has recently shown that ELF6 competes with the chromatin remodeler INO80 for chromatin binding at the transcription start sites of cell cycle gene, to precisely regulate gene expression during the cell cycle (Wang et al., 2025). Thus, it is possible to speculate that MtPKDM9B could participate in H3K27me3 demethylation of cell cycle genes to restrict cell proliferation during root development.

In conclusion, this study presents an interesting link between the regulatory mechanisms of alternative splicing, translational regulation and the removal of epigenetic repressive marks not only in the establishment of the symbiotic association between legumes and rhizobia, but also in the control of root architecture, which represents a step forward to elucidate the roles of histone demethylases in these agronomically important processes.

## Materials and Methods

### Biological material

*Medicago truncatula* Jemalong A17 seeds were obtained from the Institut national de recherche pour l’agriculture, l’alimentation et l’environnement (INRAE), Montpellier, France. *Escherichia coli* strain Top 10 or DH5a were used for vectors transformation. *Agrobacterium rhizogenes* strain Arqua1 was used for hairy roots transformation (Quandt, 1993). *Sinorhizobium meliloti* strain 1021 (Meade, 1977) or the same strain expressing RFP (Tian et al., 2012) were used as previously described (Hobecker et al., 2017).

### Constructs for plant transformation

The GATEWAY cloning system was used in this work for all constructs. *MtPKDM9B RNAi* construct was generated by the amplification of a fragment of *MtKPKDM9B* on the exon 4 using cDNA from *M. truncatula* roots as a template and primers listed in Supplemental Table 9. The *GUS RNAi* construct was generated by amplification of a *GUS* fragment using the pKGWFS7.0 vector (Karimi et al., 2002) as template and the GUS RNAi Fw and GUS RNAi Rv primers listed in Supplemental Table 9. These amplified fragments were cloned into the pENTR/D-TOPO vector (Thermo Fisher Scientific) and recombined into the destination vector pK7GWIWG2D (II) (Karimi et al., 2007). The amiR *MtPKDM9B* construct was generated following the protocol previously described (Schwab et al., 2005) by overlapping PCR using miR319 precursor into the plasmid pRS300 as PCR template to change the miR319 sequence by 21 nucleotides of *MtPKDM9B* exon 3. A first round amplifies precursor fragments with different primers combinations: A and IV miR*a, III miR*s and II miR-a, I miR-s and B (A and B primers are based on the template plasmid sequence and the rest have the 21 nucleotides of *MtPKDM9b* exon 3 mutated) using the pRS300-miR319 as template. Oligonucleotide and stem loop sequences can be found in Supplemental Table 9. These fragments were subsequently fused by PCR using A and B primers (Schwab et al., 2005). amiR MtPKDM9B was cloned into the pENTR/D-TOPO vector and then recombined into the destination vector pK7WG2D,1 (Karimi et al., 2002). The EV pK7WG2D,1 was used as a control. The pK7GWIWG2D (II) and pK7WG2D,1 vectors carrying the EgfpER as a screenable marker for early visualization and selection of the transgenic roots, were introduced into *Agrobacterium rhizogenes* ARqua1 by electroporation.

### Plant growth conditions, hairy root transformation and *rhizobium* inoculation

Seeds were surface sterilized and germinated on 10% (w/v) agar water plates at 25°C in the dark for 24 h. Transgenic roots were generated by *A. rhizogenes-*mediated transformation as described previously (Boisson-Dernier et al., 2001) and transferred to Petri dishes containing agar Fahraeus media supplemented with 8mM KNO_3_ and 12.5 µg/ ml of Kanamycin for 14 days. Seedlings were grown at 25°C with a 16/8-h day/night cycle with photosynthetically active radiation of 200 μmol.m^−2^.s^−1^ using mixed lighting containing four OSRAM cool daylight L36W/765 tubes per one OSRAM FLUORA L36W/77 tube. For root architecture analysis transgenic plants were transferred to slanted 12 cm square petri dishes containing Fahraeus media supplemented with 8mM KNO_3_ or not and grown under the conditions described above for 14 or 21 days. For inoculation with rhizobia, plants that developed hairy roots were transferred to slanted boxes containing Fahraeus media free of nitrogen covered with sterile filter paper. Roots were inoculated with 10 mL of a 1:1000 dilution of *S. meliloti* 1021 (Meade, 1977) or the same strain expressing RFP (Tian et al., 2012) culture grown in liquid TY media until OD_600_ reached 0.8, or with 10 mL of water as a control (mock treatment). One hour later, the excess liquid was discarded, and seedlings were incubated vertically under the growth conditions described above.

### Classification and phylogenetic analysis of demethylases

The sequence of proteins with JMJ were extracted from the *Medicago truncatula* genome database (https://medicago.toulouse.inra.fr/MtrunA17r5.0-ANR/) and the TAIR database (https://www.arabidopsis.org). The alignment of multiple sequences and phylogenetic tree was constructed in MEGA-X software using the Neighbor Joining (NJ) method, with 10000 bootstrap replicates.

### RNA Extractions and RT-qPCR

Root tissue was harvested using frozen with liquid N_2_ and stored at -80°C. Total RNA extraction was performed using harvested root tissue with Trizol according to the manufacturer’s instructions (Thermo Fisher). RNA concentration was determined by measuring A260 using a Nanodrop ND-1000 (Nanodrop Technologies), and RNA integrity was evaluated by electrophoresis on 1% (w/v) agarose gels. Total RNA was treated with DNase I (Promega) and was subjected to first-strand cDNA synthesis using MMLV reverse transcriptase (Promega). Expression analysis by RT-qPCR was performed using the iQ SBR Green Supermix kit (BioRad) and the CFX96 qPCR system (BioRad) as previously described (Reynoso et al., 2013). For each pair of primers, the presence of a unique product of the expected size was verified on 1.2% (w/v) agarose gels. In all cases, negative controls without template or without reverse transcription were included. Expression values were normalized to *HIS3L*, a reference transcript for normalization of RT-qPCR data previously reported in a geNORM analysis (Reynoso et al., 2013).

### Phenotypic Analyses

For nodulation analysis, nodule number was recorded at different time points after inoculation with *S. meliloti* as described previously by Hobecker et al. (2017). Nodule size was measured from digital pictures using ImageJ at 21 dpi using more than 30 nodules per construct. Nodules and infection events were quantified in at least 50 independent roots per construct inoculated with a *S. meliloti* strain expressing RFP (Tian et al., 2012). Microcolonies and infections threads were visualized, quantified, and imaged at 7 dpi with a *S. meliloti* strain expressing RFP on roots transformed using an IX51 inverted microscope (Olympus). For SYTO9/propidium iodide staining, nodules were excised and embedded in 6% (w/v) low melting agarose. Nodule sections of 60 μm were obtained using a VT1000 S vibratome (Leica Microsystems). Staining was performed using the LIVE/DEAD™ *Bac*Light™ Bacterial Viability kit (Invitrogen). Images of nodule sections were acquired with an inverted SP5 confocal microscope (Leica Microsystems) and IX51 inverted microscope (Olympus). Images were processed with the LAS Image Analysis software (LeicaMicrosystems) and ImageJ. For root architecture analysis, the length of primary and lateral roots, and the number of lateral roots per centimeter of primary root were determined at 14 and 21 days after transplanting to slanted plates containing agar-Fahraeus medium supplemented with 8 mM KNO3 or without nitrogen. Roots emerged directly from the sectioned radicle transformed with *A. rhizogenes* were considered primary roots in the hairy root system and roots that emerged from these primary roots were considered lateral roots. Three biological replicates were performed for each experiment. The statistical significance of the differences for each parameter was determined by unpaired two-tailed Student’s t tests for each construct.

### ChIP-seq and ChIP-PCR assay

ChIP assays were performed on transgenic roots of *MtPKDM9B* RNAi and *GUS* RNAi 48 hpi with *S. meliloti* (Sm) or water (*mock*) using a commercial anti-H3K27me3 antibody (Diagenode) essentially as previously described (Kirolinko et al., 2024). Three grams of root tissue was cross-linked in 1% (v/v) formaldehyde for 15 min under vacuum. Crosslinking was quenched with 1.25M glycine and vacuum. The cross-linked tissue was ground and filtered with 100 µm filter then nuclei were isolated with a sucrose cushion and lysed in Nuclei Lysis Buffer (1% SDS, 50mM Tris-HCl pH 8, 10 mM EDTA pH 8). Cross-linked chromatin was sonicated using a Bioruptor Plus sonicator (Diagenode) (30 sec on/30 sec off pulses; 30 times). An aliquot of 10% of this was taken to use as input. The complexes were immunoprecipitated with anti-H3K27me3 antibodies or anti-IgG antibody (as a negative control), overnight at 4°C with gentle shaking, and incubated for 1 h at 4°C with 50 mL of Protein G (Thermo Fisher). The beads were washed 2 times for 5 min in ChIP Wash Buffer 1 (0.1% (w/v) SDS, 1% (v/v) Triton X-100, 20 mM Tris-HCl pH 8, 2 mM EDTA pH 8, 150 mM NaCl), 2 times 5 min in ChIP Wash Buffer 2 (0.1% (w/v) SDS, 1% (v/v) Triton X-100, 20 mM Tris-HCl pH 8, 2 mM EDTA pH 8, 500 mM NaCl), 2 times for 5 min in ChIP Wash Buffer 3 (0.25 M LiCl, 1% (v/v) NP-40, 1% (w/v) sodium deoxycholate, 10 mM Tris-HCl pH 8, 1 mM EDTA pH 8) and twice in TE (10 mM Tris-HCl pH 8, 1 mM EDTA pH 8). Reverse-crosslinking was performed in ChIP DNA and inputs with PK buffer (Tris-Hcl10mM, EDTA, NaCl 50mM) and 20mg/ml proteinase K incubated 4h at 65°C. Reverse-cross-linked DNA was extracted with phenol-chloroform. DNA was precipitated with 1 µl of glycogen, 50 µl AcNa (PH=5,2) and 2 volumes of ethanol with centrifugation. The precipitated DNA was resuspended in 30 µl of nuclease-free water and added 0.5 µl of RNAse A. In ChIP-qPCR experiments, enrichment was calculated as the ChIP H3K27me3/input percent normalized by the ChIP IgG/input. ChIP or input DNA was used for ChIP-Seq library construction using *Trans*NGS Tn5 Library Prep Kit for Illumina (Transgen Biotech) and AMPure XP beads for DNA and library purification (Beckman Coulter Genomics). The quality of the libraries was assessed with Agilent 2100 Bioanalyzer (Agilent) and the concentration was determinate by qPCR. Libraries were sequenced on an Illumina NovaSeq platform using paired end reads (2 × 150 bp) at Novogene (https://www.novogene.com/us-en/). After sequencing, an average of 51 million reads were obtained for each ChIP library, with nearly 17% of the reads mapping at a single locus in the *M. truncatula* genome 5.0, whereas only 2% of the reads obtained in the sample immunoprecipitated with α-rabbit IgGs mapped to a single locus.

### ChIP-seq Computational analysis

Paired-end sequencing was performed. Reads were quality controlled using FASTQC with filter parameters quality cutoff -30 and cut nextera adapter (https://usegalaxy.org/) ChIP and inputs-seq reads were aligned to *M. truncatula* genome v5 using Bowtie (Langmead, 2010). The significantly enriched regions were identified using MACS2 (Zhang et al., 2008). For peak annotation ChIPseeker was used (Yu G et al.,2015). Integrative Genomic Viewer (IGV) (Thorvaldsdottir et al., 2013) was used to visualize the read with a coverage bigwig file generated by bamCoverage tool (Ramírez et al., 2016). Regions differentially enriched in H3K27me3 between samples were identified using DIFFREPS (Shen et al., 2013) with parameters of p-value 0,05; z-score cutoff 2; G-test; windows 1000. R studio was used to filter the tables for p-value 0,05; |log_2_ fold change| > 1, and for peak annotation such as proximity to genes and overlapping on genomic features.

### RNA-seq sample preparation and sequencing

Toral RNA was obtained from two biological replicates of *MtPKDM9B* RNAi and *GUS* RNAi transgenic roots 48 hpi with *S. meliloti* (Sm) or water (mock) using Trizol. RNA concentration was determined by measuring A260 using Qubit (Thermo Fisher) according manufacturer’s instructions and analyzed on Agilent Bioanalyzer as previously described (Traubenik et al., 2020). Illumina compatible RNA-seq libraries were prepared using the YourSeq Duet Full Transcript & 3’-Digital Gene Expression (FT & 3’-DGE) RNAseq Library Kit following manufacturer’s instructions (Amaryllis Nucleics). Libraries were sequenced on an Illumina NovaSeq platform using paired end reads (2 × 150 bp) at Novogene (https://www.novogene.com/us-en/). An average of 62 million reads per library were obtained with more than 88% of bases with quality scores ≥30.

### RNA-seq computational analysis

Raw sequencing reads were subjected to quality control using FastQC. Adapter sequences and low-quality bases were removed using Cutadapt in a Galaxy environment (https://usegalaxy.eu/). Filtered reads were aligned to the *M. truncatula* reference genome V.5 available at MtrunA17r5.0-ANR using HISAT2 (Kim et al., 2019). Mapping efficiency was high with 94% of reads mapping to the reference genome and ∼75% of reads mapping uniquely. Read counts were generated using featureCounts considering only uniquely mapped reads. Differential gene expression analysis between experimental conditions was performed using DESeq2 (Love et al., 2014) within the Galaxy environment. Log_2_ fold changes and significance values. Genes with an adjusted p-value < 0.05 and a |log_2_ fold change| > 1 were considered significantly differentially expressed. Principal component analysis (PCA), sample distance clustering, and visualization of differential expression patterns (heatmaps and volcano plots) were conducted using DESeq2.

## Data availability

The ChIP-seq and RNA-seq raw data, as well as processed data, generated here were deposited at GEO under accession number GSE318381 and GSE337466, respectively.

## Funding

This work was supported by grants of the Agencia Nacional de Promoción de la Investigación, el Desarrollo Tecnológico y la Innovación (Agencia I+D+I) of Argentina, FONCYT (PICT2019-00554, PICT2019-01970 and PICT-2021-I-A-00170), the Ministerio de Ciencia, Tecnología e Innovación (MINCyT) of Argentina (RIBOLEG, CONVE-2023-100766842), M.A.R., F.A.B., and M.E.Z. are members of CONICET, Argentina. M.F., M.Y., EI are funded by a CONICET fellowship.

## Author contributions

M.F., M.E.Z., F.B., and M.A.R. designed the research. M.F., S.T., M.Y., and E.I. performed the research. M.F. and M.E.Z. wrote the original draft of the article. M.F., S.T., M.A.R., F.B. and M.E.Z. analyzed the data. M.F., S.T., E.I.; M.A.R., F.B., and M.E.Z. reviewed and edited the article. Funding was acquired by M.A.R; F.B. and M.E.Z. M.E.Z. and M.A.R supervised the study.

## Acknowledgements

We acknowledge Dr. Pablo Cerdán, Dr. Paula Cassatti and Dr. Hernán Rosli for helpful discussions. No conflict of interest declared.

## Supplemental Figures

**Supplemental Figure 1.**
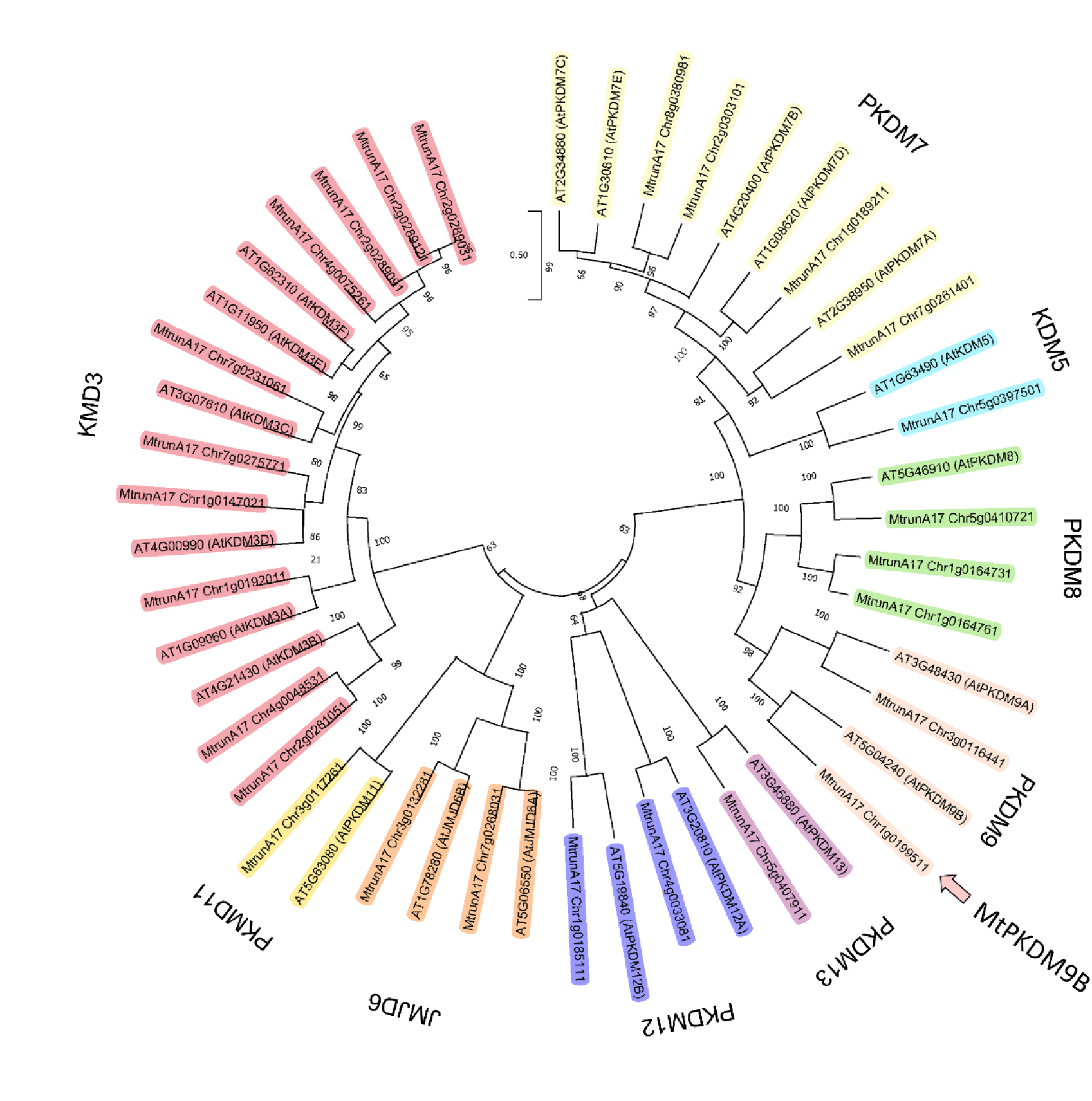
Phylogenetic tree of the JmjC domain protein family from Arabidopsis and *M. truncatula*. The tree was constructed from a multiple sequence alignment using MEGA-X and the Neighbor joining method, with 10,000 iterations. Nine groups of proteins with JmjC domains were identified, indicated by different colored shading. The pink arrow points to the MtPKDM9B protein.

**Supplemental Figure 2.**
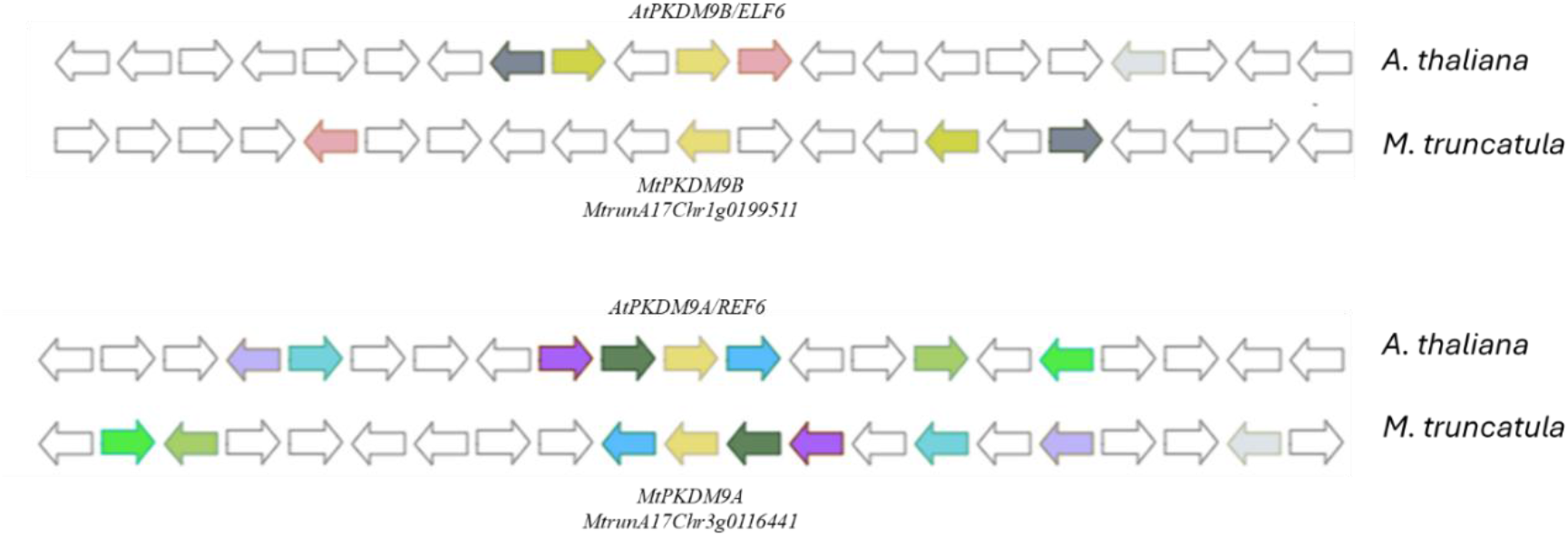
Synteny analysis. Synteny between *AtPKDM9B* and *MtPKDM9B* genes, and between *AtPKDM9A* and *MtPKDM9B* genes. Genes are indicated by arrows. PKDM9B and PKDM9A genes are indicated in light yellow arrows. Those genes belonging to the same family are indicated by the same color. Syntenic analysis was performed using Dicots PLAZA 5.0 (https://bioinformatics.psb.ugent.be/plaza.dev/_dev_instances/feedback/synteny/synteny) using default parameters.

**Supplemental Figure 3.**
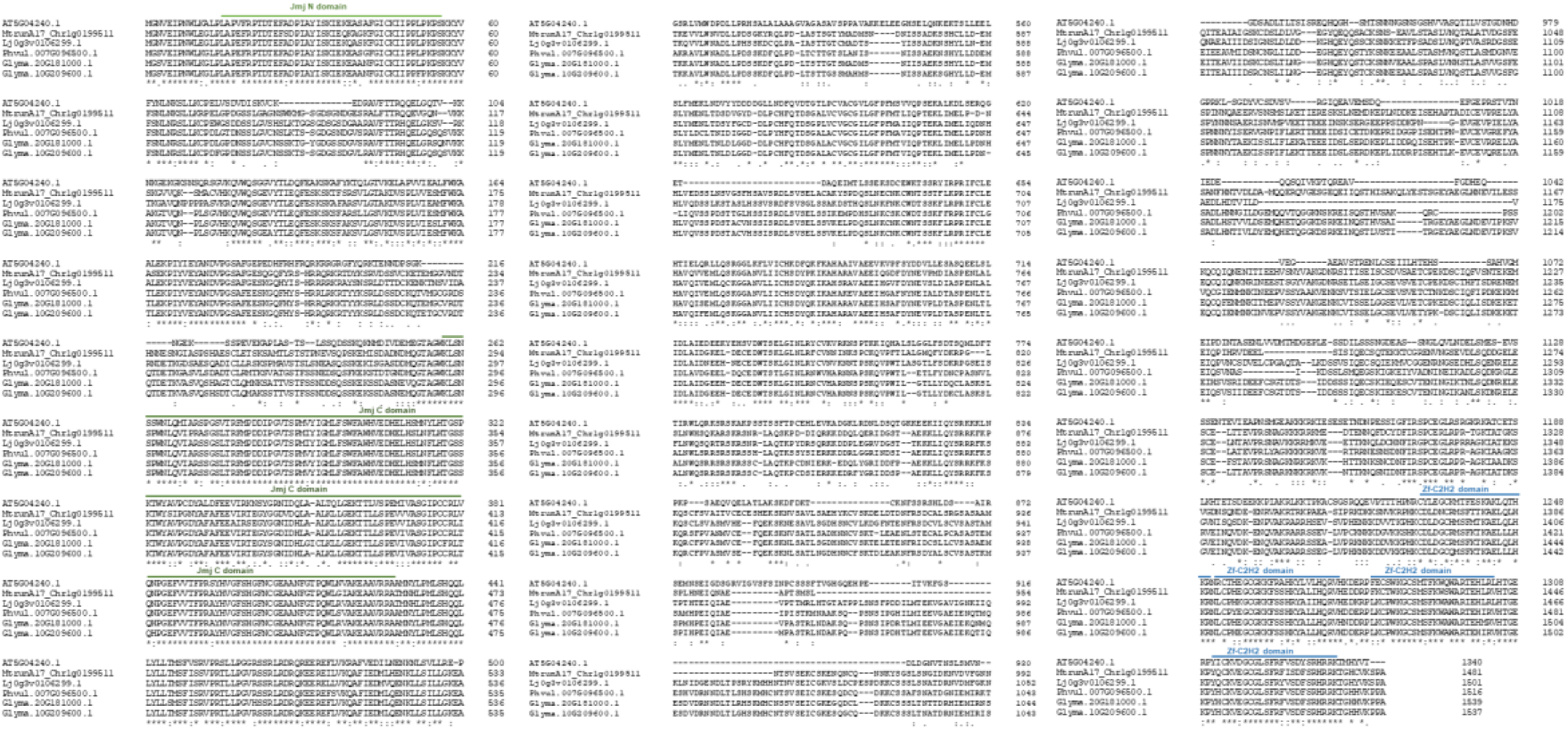
Multiple sequence alignment of PKDM9B proteins from Arabidopsis and legumes. Amino acid sequences from PKDM9B protein from Arabidopsis (At5G04240.1), *M. truncatula* (MtrunA17_Chr1g0199511), *Lotus japonicus* (Lj0g3v0106299.1), *Phaseolus vulgaris* (Phvul.007G096500.1) and *Glycine max* (Glyma.20G181000.1 and Glyma.10G209600.1) were retrieved from The Arabidopsis Information Resource (TAIR) (https://www.arabidopsis.org/), *Medicago truncatula* Jemalong A17 5.0 genome (https://medicago.toulouse.inra.fr/MtrunA17r5.0-ANR/) and Phytozome 14 (https://phytozome-next.jgi.doe.gov/) web portals. Sequences were aligned using Clustal Omega Multiple Sequence Alignment (MSA) (Madeira et al., 2024). The Jmj N-terminal (Jmj N-ter) and C-terminal (Jmj C-ter) domains are indicated by green lines and the four zinc Finger C2H2 (ZF-C2H2) domains are indicated by blue lines above the sequence alignment. Asterisk (*) Indicates identical amino acids, colon (:) indicates conservative substitution and dot (.) indicates semi-conservative substitutions.

**Supplemental Figure 4.**
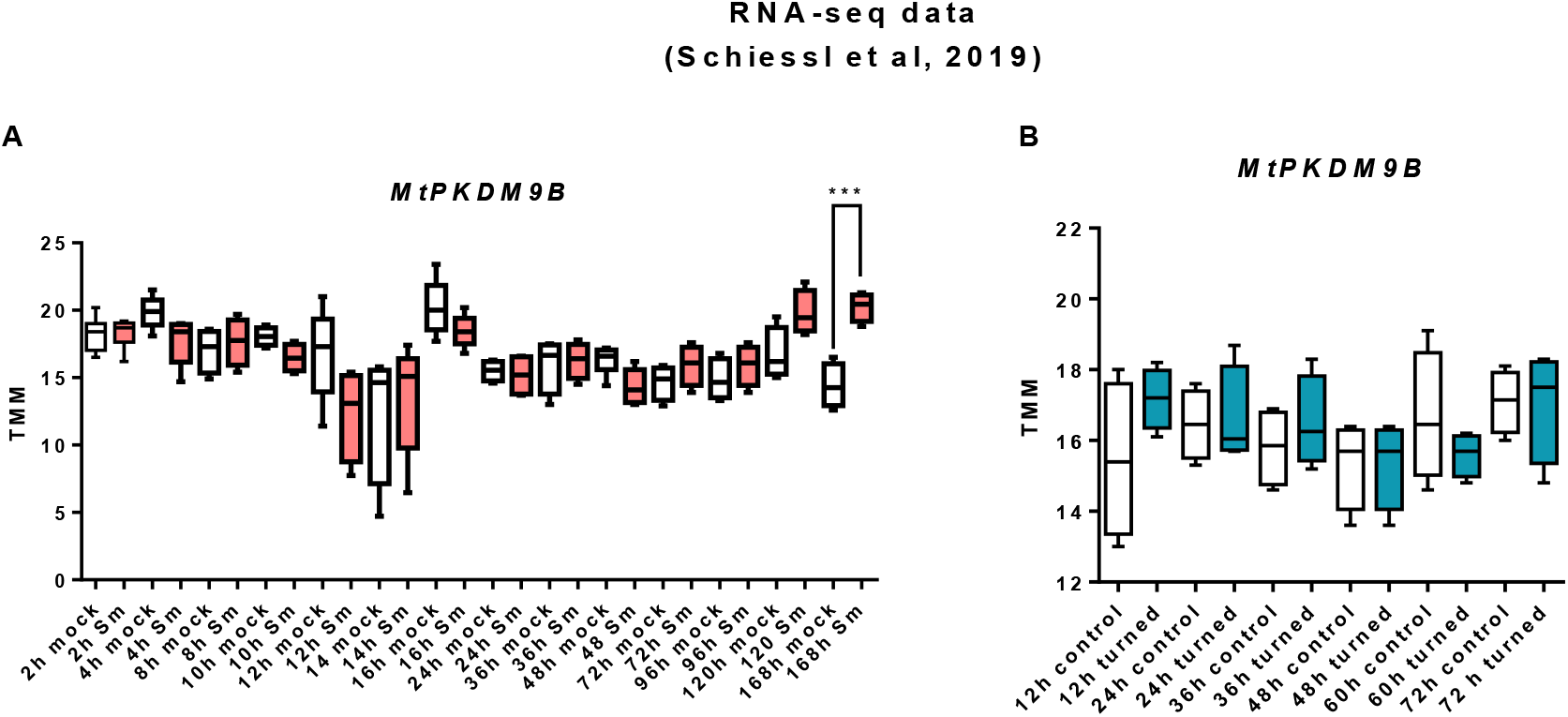
Expression analysis of *MtPKDM9B* during nodule (A) and lateral root (B) formation. *MtPKDM9B* levels in *M. truncatula* roots at indicated times point after spot inoculation with *S. meliloti* **(A)** or after induction of lateral root formation by turning seedlings 135° **(B)** were obtained by RNA-seq data reported by (Schiessl et al., 2019). Values are expressed as TMM (Trimmed mean of M-values). Asterisks indicate statistically significant differences using a Student t-test (***: p≤0.001).

**Supplemental Figure 5.**
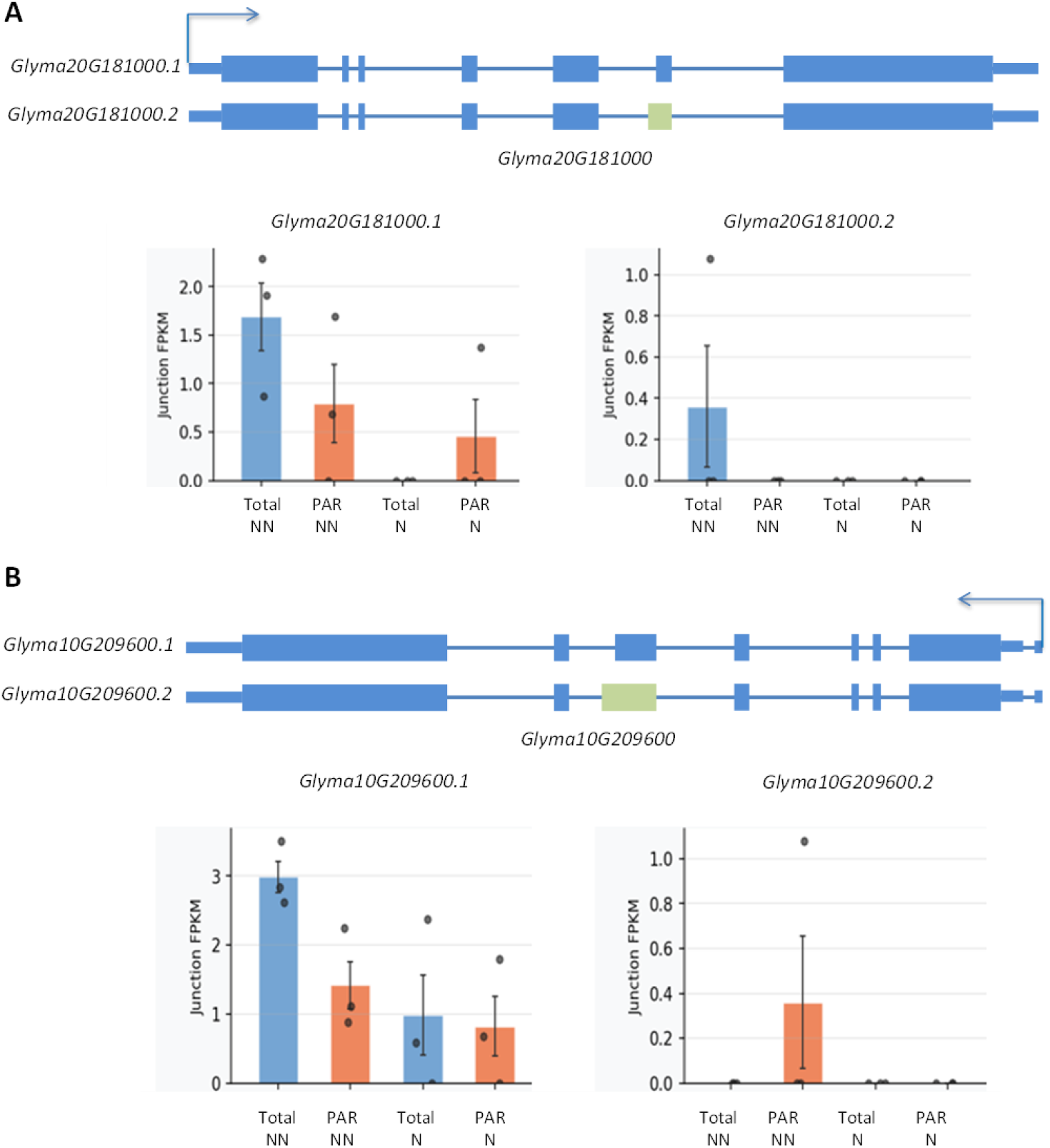
Alternative spliced variants of *Glyma20G181000* and *Glyma10G209600* and their expression and association with polyribosomes during nodulation. **(A)** *Glyma20G181000.1 and Glyma20G181000.2* variants produced due to an alternative 3’acceptor site. Abundance of *Glyma20G181000.1 and Glyma20G181000.2* in Total RNA (blue) and Polyribosomes Associated mRNA (PAR, orange) samples from non nodulated (NN) and nodulated roots (N) *Glyma20G181000.1*, but not the alternative transcript *Glyma20G181000.2* was associated with translating polyribosomes. (**B**) *Glyma10G209600.1* and *Glyma10G209600.*2 variants produced due to an alternative 5’donnor site. *Glyma10G209600.1* encodes the full-length protein, whereas *Glyma10G209600.*2 contains a premature stop codon. Abundance of *Glyma10G209600.1* and *Glyma10G209600.*2 in Total (blue) and PAR (orange) samples from non NN and N roots. In NN both variants were found to be associated with translating polyribosomes. In NN roots the alternative variant *Glyma.10G209600.2* is absent in translating poyribosomes, whereas the canonical splice variant *Glyma.10G209600.1* remains associated with translating polyribosomes. RNA sequencing data was obtained from Sainz et al., (2022).

**Supplemental Figure 6.**
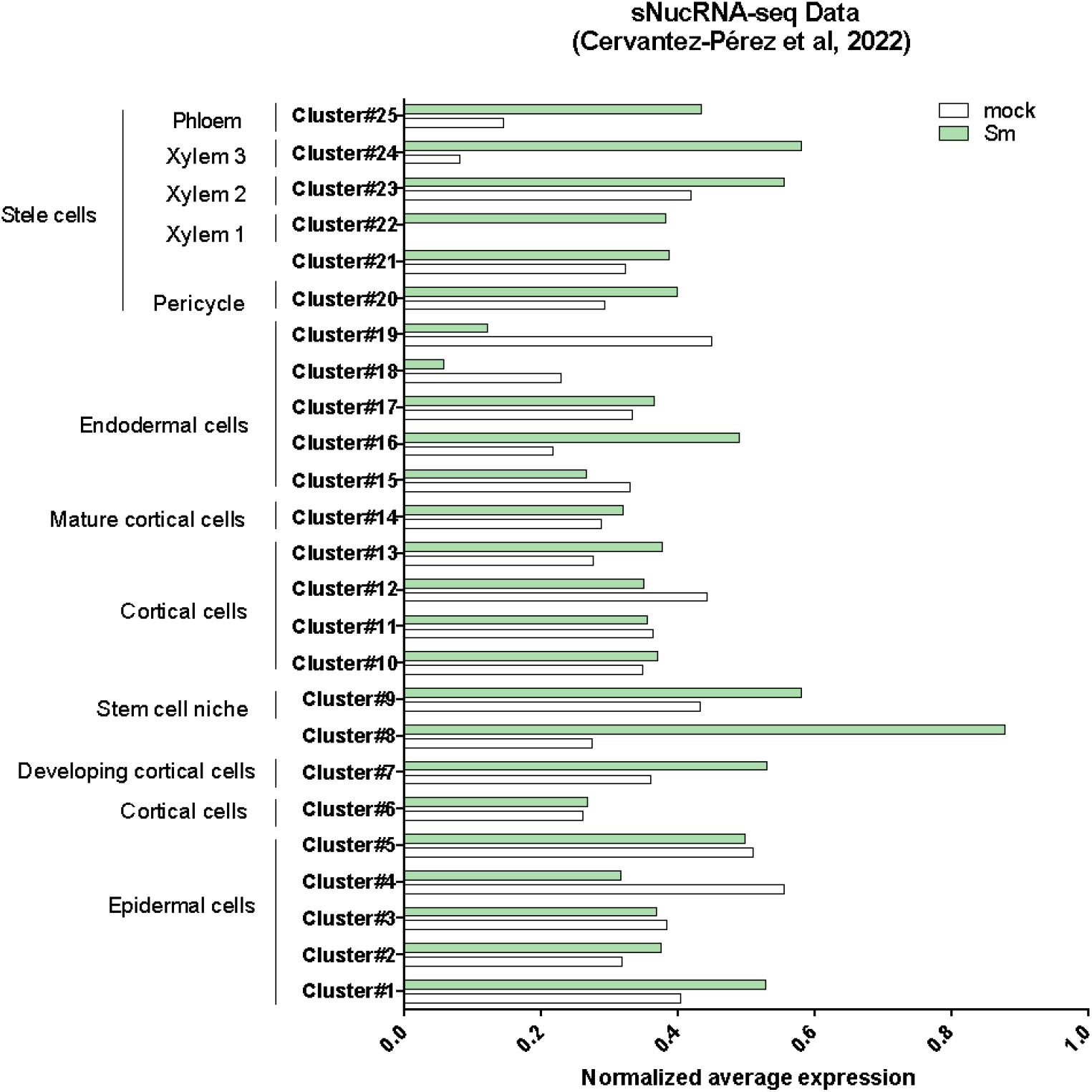
M*t*PKDM9B normalized transcript levels in clusters of individual nuclei in response to the inoculation with *S. meliloti*. *MtPKDM9B* expression levels obtained from single-nuclei RNA-seq data from mock and *S. meliloti* roots at 48 hpi (Cervantes-Perez and Libault, 2022) separated into 25 clusters corresponding to different cell types.

**Supplemental Figure 7.**
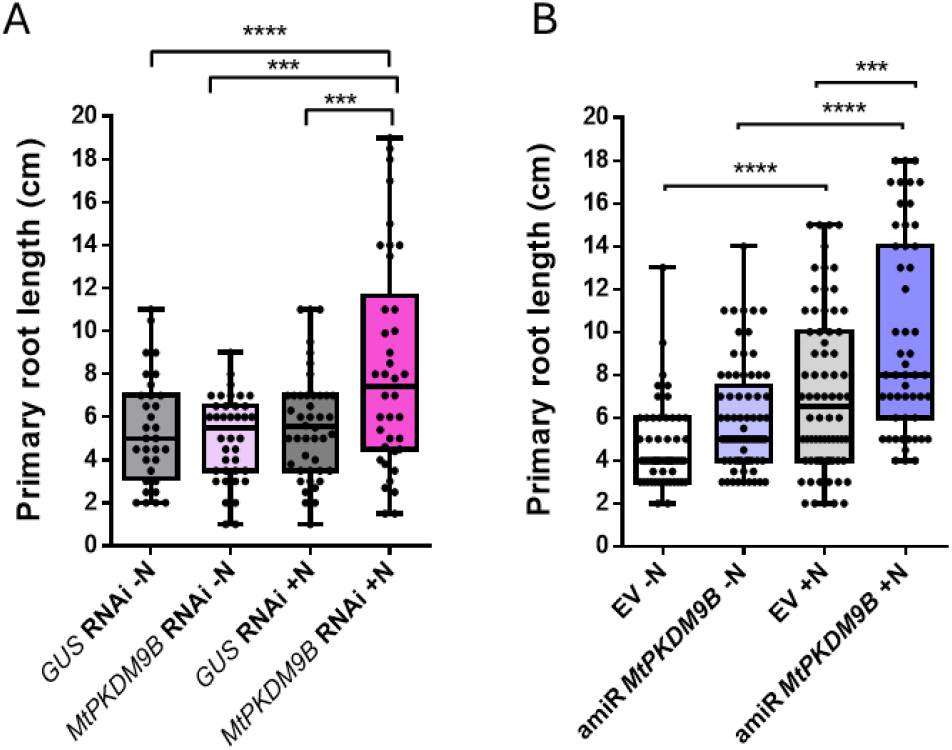
Silencing of *MtPKDM9B* promotes primary root growth at 21 days after transplantation. Primary root length of *MtPKDM9B* RNAi **(A)** and amiR *MtPKDM9B* **(B)** and their controls at 21 days after transplantation plants to media with free of nitrogen (-N) or supplemented with KNO_3_ (+N). Boxes extend from the 25th to 75th percentiles, the middle line is the median, and whiskers extend to the minimum and maximum values of three technical replicates with at least 35 plants each. Asterisks denote statistically significant differences in an unpaired two-tailed Student’s t-test (∗∗∗p ≤ 0.001 and ****p ≤ 0.0001).

**Supplemental Figure 8.**
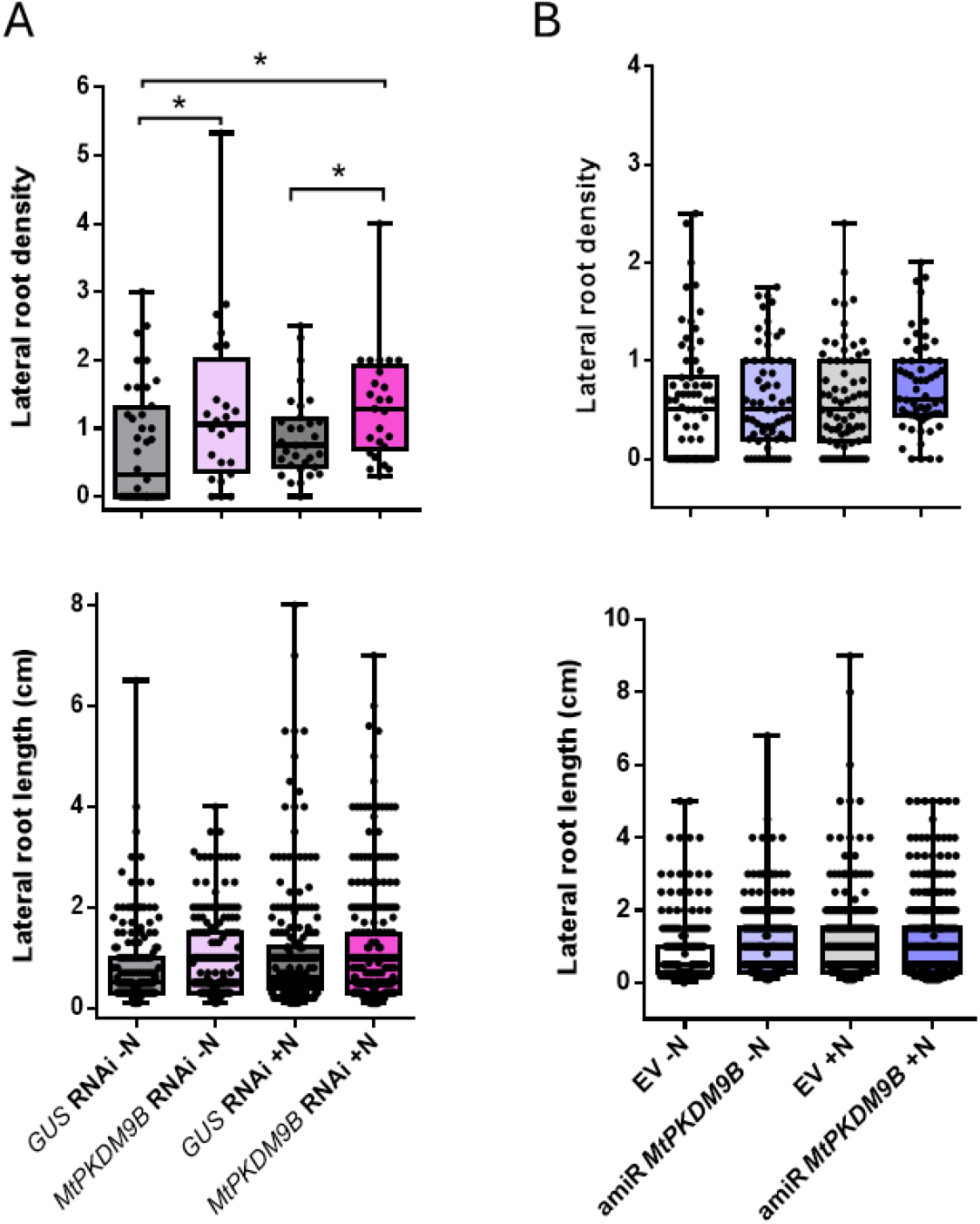
Silencing of both *MtPKDM9B.1 and MtPKDM9B.2* variants alters lateral root density. Lateral root density and length measured in *MtPKDM9B* RNAi roots **(A)** and amiR *MtPKDM9B* **(B)** and their control *GUS* RNAi and EV, respectively, at 14 days after transplantation of plants to media free of nitrogen (- N) or supplemented with KNO_3_ (+N). Boxes extend from the 25th to 75th percentiles, the middle line is the median, and whiskers extend to the minimum and maximum values of three technical replicates with at least 35 plants each. The asterisk denotes statistically significant differences in an unpaired two-tailed Student’s t-test with p ≤ 0.05.

**Supplemental Figure 9.**
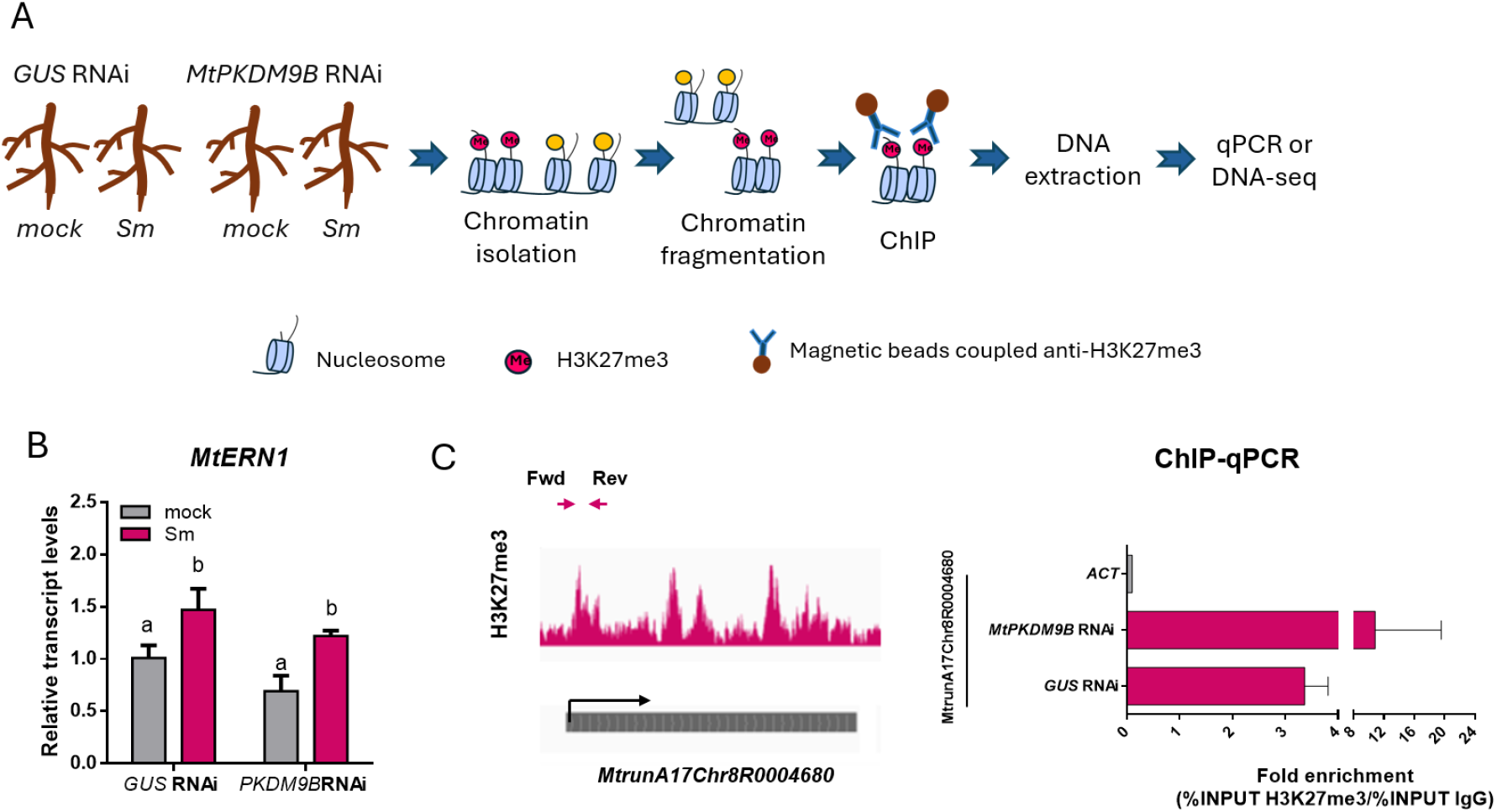
ChIP experimental design and validation. **(A)** Experimental design for the ChIP experiment. Tissue from *MtPKDM9B* RNAi and *GUS* RNAi roots at 48 hpi with water (mock) or *S. meliloti* (Sm) were harvested. Nuclei were purified and chromatin was isolated and fragmented to 150 pb by sonication. Samples were immunopurified using a commercial α-H3K27me3 antibody or an α-rabbit IgG antibody as a control. DNA was extracted from the Input and ChIP samples. Purified DNA was used for qPCR or for construction of DNA libraries for Illumina sequencing. **(B)** Verification of induction of the early symbiotic marker *MtERN1* in *MtPKDM9B* RNAi and *GUS* RNAI roots at 48 hpi with water (mock) or *S. meliloti* (Sm). Values are the mean and standard error of the mean. Different letters indicate that values are significantly different in an unpaired two-tailed Student’s t-test with p ≤ 0.05. **(C)** Genome browser capture of the locus *MtrunA17Chr8R0004680*, which exhibits high levels of H3K27me3 in root tissues (left panel). Primers flanking the region to be amplified by qPCR are indicated with purple arrows. Black arrow indicates the transcriptional start site. Data was visualized in the Integrative Genomic Viewer (IGV) using ChIP-seq data antibody from root tissues previously generated using α-H3K27me3 in roots by Pecrix et al., (2018) and in root tip . ChIP-qPCR from *MtPKDM9B* RNAi and *GUS* RNAI root tissue. Fold enrichment was estimated as the percentage of the Input in the ChIP-H3K27me3 sample relative to the percentage of the Input in the ChIP IgG sample. Amplification of the *ACTIN11* (*ACT*) loci, which contain low H3K27me3 levels, was used as a negative control.

**Supplemental Figure 10.**
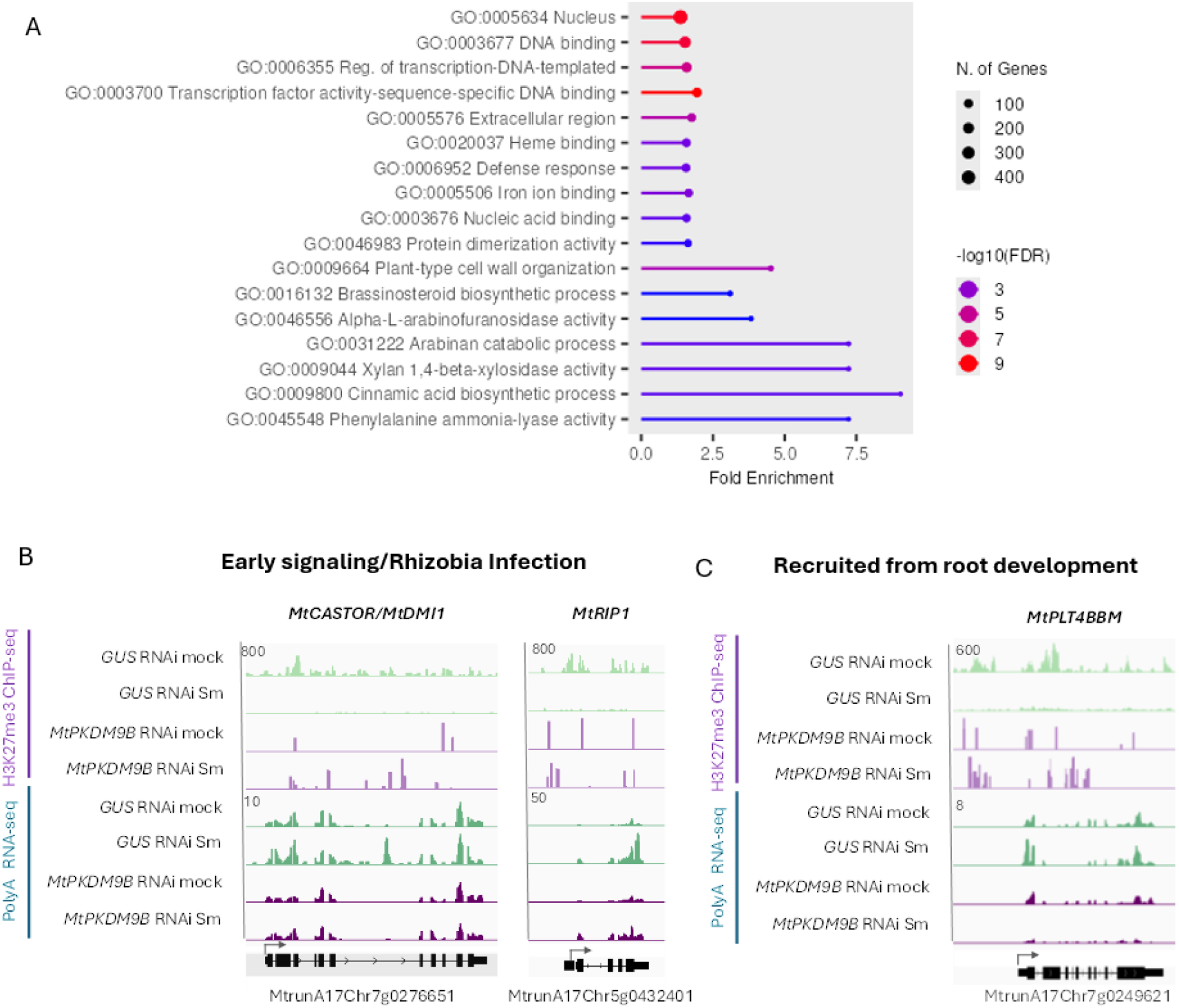
Differentially H3K27me3-methylated regions (DHMRs) in putative targets of *MtPKDM9B*. **(A)** Gene ontology analysis of hypermethylated DHMRs detected in the comparison between *MtPKDM9B* RNAi versus *GUS* RNAi samples under *S. meliloti* inoculated conditions. Numbers of genes in each category are presented by the size of the circles and the probability as -log_10_ of fold discovery rate (FDR) is represented by different colors. GO categories are presented from top to bottom based on the number of genes in each category. GO analysis was performed with ShinyGO 0.85.1 available at https://bioinformatics.sdstate.edu/go/ (Ge et al., 2019). **(B-C)** Integrative Genomic Views (IGV) of ChIP-seq signals with H3K27me3 and RNA-seq reads for genes involved in early signaling/rhizobial infection, *MtCASTOR* and *MtRIP1* **(B)**, and a transcription factors that function in root nodule symbiosis that have been recruited from root developmental program *MtPLT4/BBM* **(C)** in mock and Sm inoculated *GUS* RNAi (green) and *MtPKDM9B* RNAi (purple) root samples. Normalized reads are presented. Gene models are presented below, and arrows indicate the transcriptional star site (TSS). Numbers on the top left indicate the maximum read value of the scale, which was the same for each gene in all samples.

**Supplemental Figure 11.**
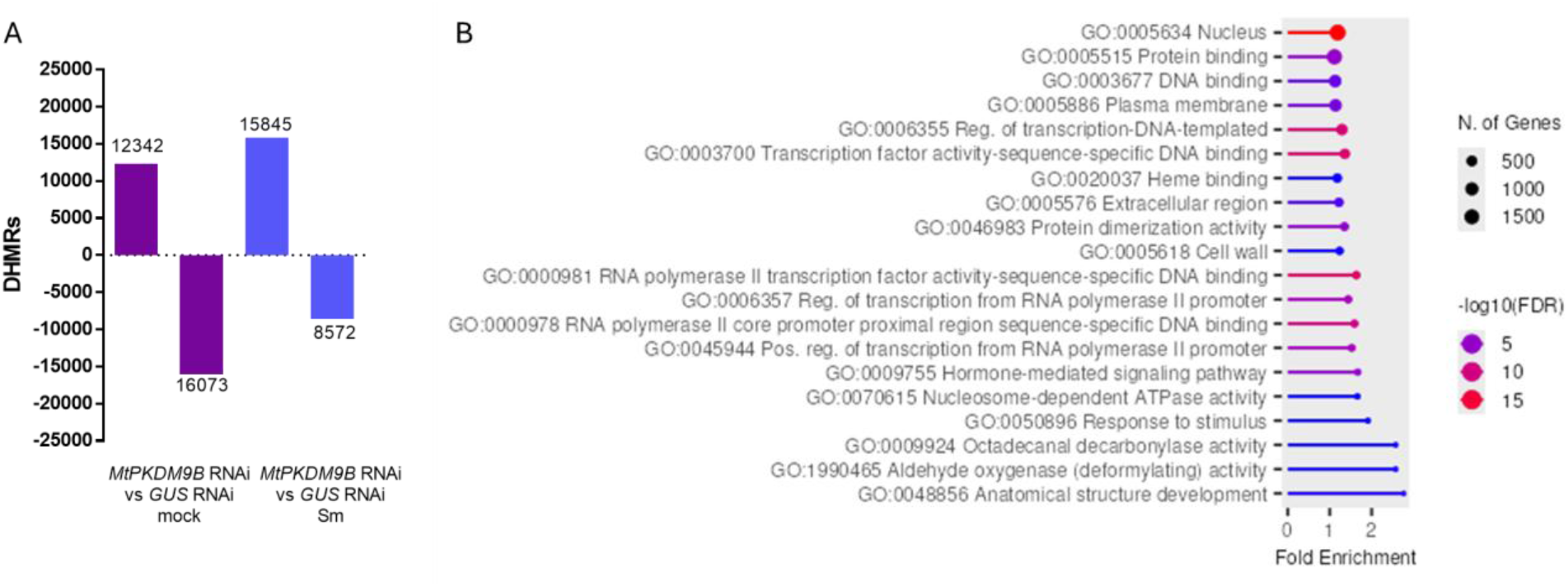
Differentially H3K27me3-methylated regions (DHMRs) between *GUS* RNAi and *MtPKDM9B* RNAi samples under mock and rhizobia inoculated conditions. **(A)** DHMRs were identified by comparing ChIP-seq data in the *GUS* RNAi and *MtPKDM9B* RNAi samples under mock and *S. meliloti* (Sm) inoculated conditions using the *diffreps* tool with a 1000-bp window, a 1<Log_2_FC<-1 and a p-value< 0.05. Positive and negative bars represent hyper- and hypo-methylated DHMRs, respectively. **(B)** Gene ontology analysis of hypermethylated DHMRs detected in the comparison between *GUS* RNAi and *MtPKDM9B* RNAi samples under *S. meliloti* inoculated conditions. Numbers of genes in each category are presented by the size of the circles and the probability as -log_10_ of fold discovery rate (FDR) is represented by different colors. GO categories are presented from top to bottom based on the number of genes in each category. GO analysis was performed with ShinyGO 0.85.1 available at https://bioinformatics.sdstate.edu/go/ (Ge et al., 2019).

**Supplemental Figure 12.**
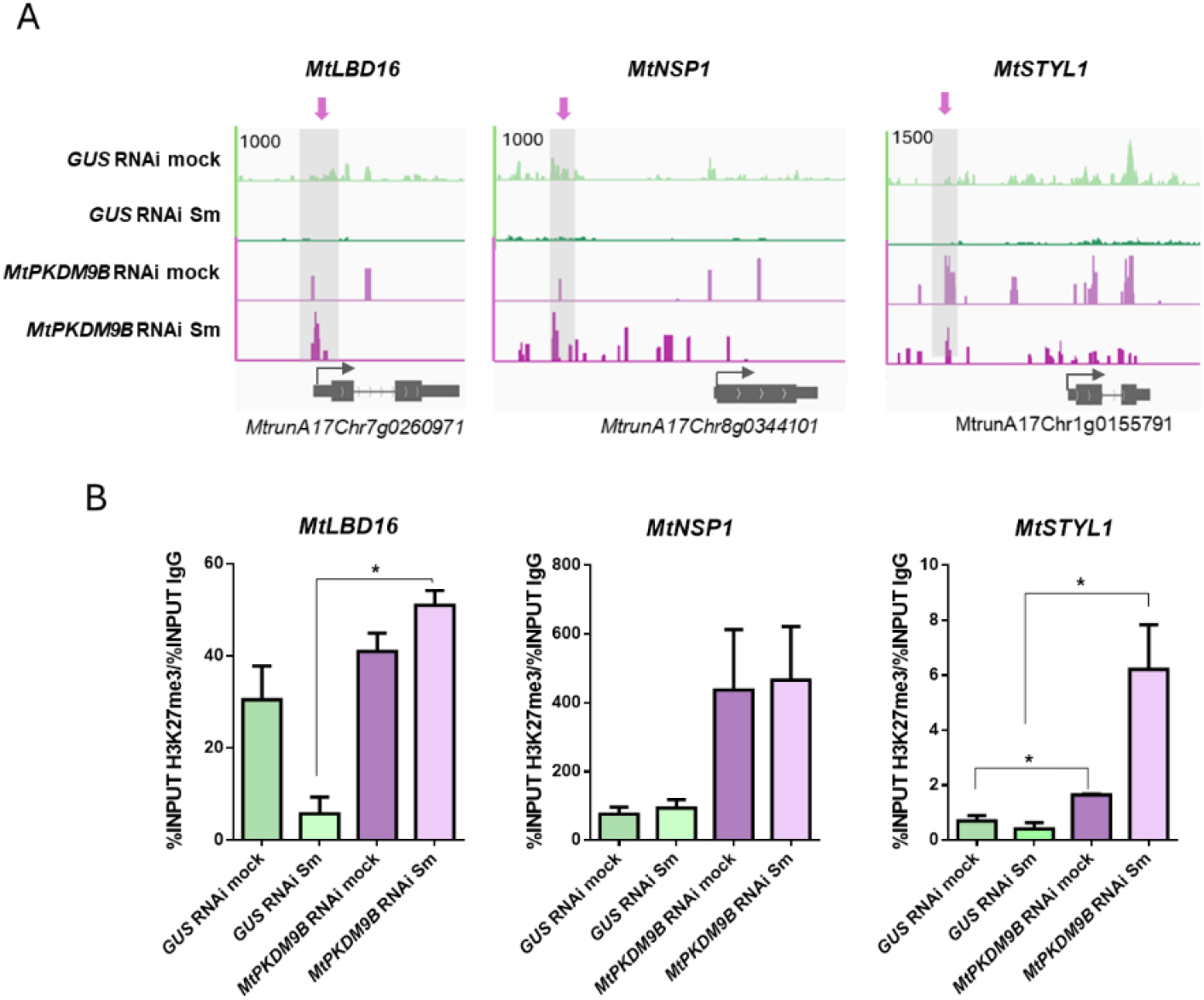
Knockdown of *MtPKDM9B* affects methylation of genes required for bacterial infection and nodule development. **(A)** Integrative Genomic Views (IGV) captures of ChIP-seq signals with H3K27me3 for *MtLBD16*, *MtNSP1* and *MtSTYL1* in mock and *S. meliloti* (Sm) inoculated *GUS* RNAi (green) and *MtPKDM9B* RNAi (purple) root samples. Normalized reads are presented. Shade boxes are DHMRs that are hypomethylated in Sm samples as compared to mock samples for each gene. Gene models are presented below, and arrows indicate the transcriptional star site (TSS). Numbers on the top left indicate the maximum read value of the scale, which was the same for each gene in all samples. **(B)** Validation of ChIP-seq data by ChIP-qPCR performed for *MtLBD16*, *MtNSP1* and *MtSTYL1* on ChIP samples with H3K27me3 in *GUS* RNAi (green) and *MtPKDM9B* RNAi (purple) samples at 48 hpi with water (mock) or *S. meliloti* (Sm) using primers flanking the DHMRs for each gene (Supplemental Table 9). Fold enrichment was estimated as the percentage of the Input in the ChIP-H3K27me3 sample relative to the percentage of the Input in the ChIP IgG sample. The asterisk indicates statistically significant differences in a two-tailed unpaired Student’s t-test (∗p ≤ 0.05)

**Supplemental Figure 13.**
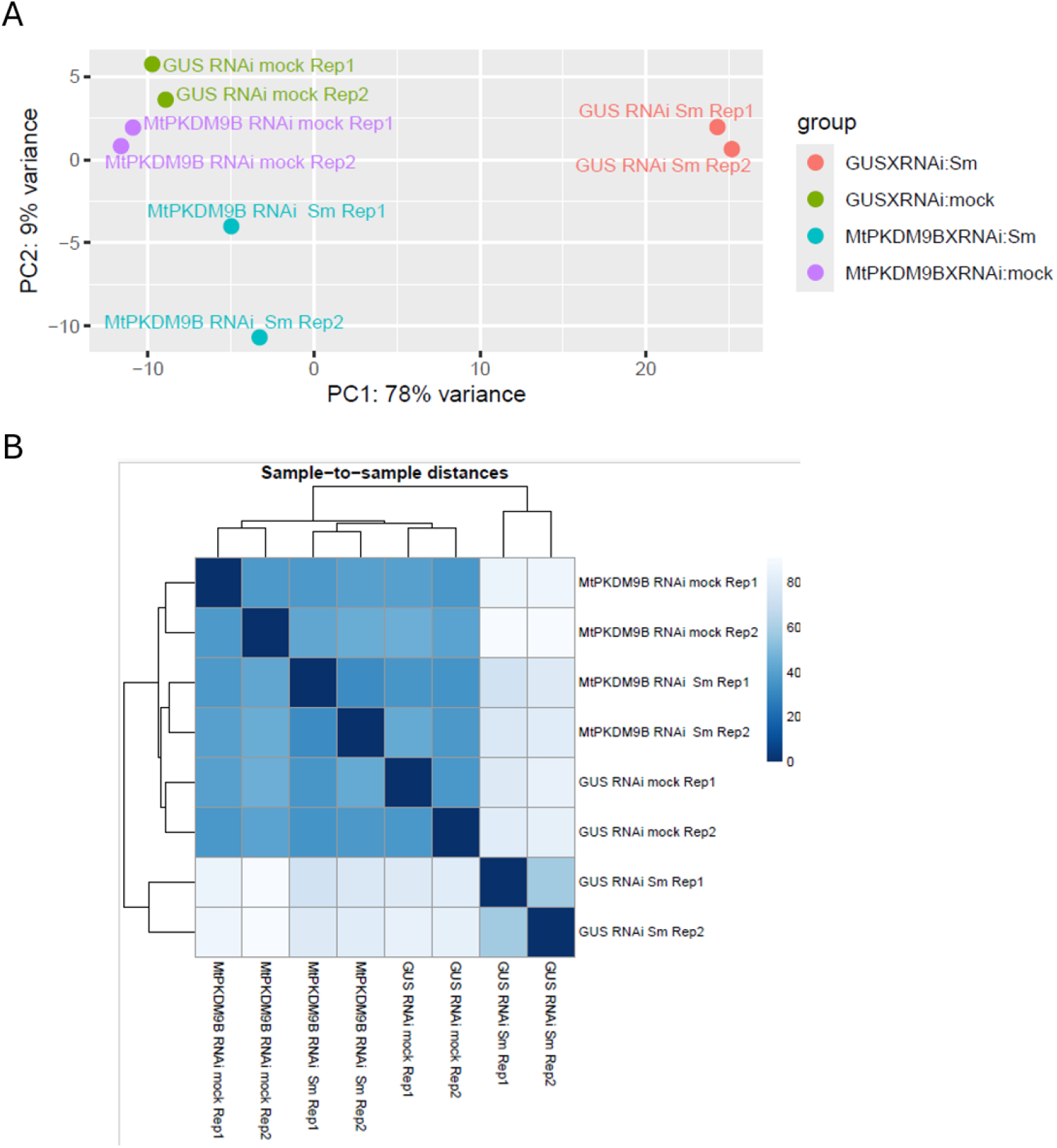
Reproducibility of RNA-seq samples. **(A)** Principal Component Analysis (PCA) and **(B)** Sample-to sample distance of mock and *S. meliloti* inoculated (Sm) *GUS* RNAI and *MtPKDM9* RNAi root samples.

**Supplemental Figure 14.**
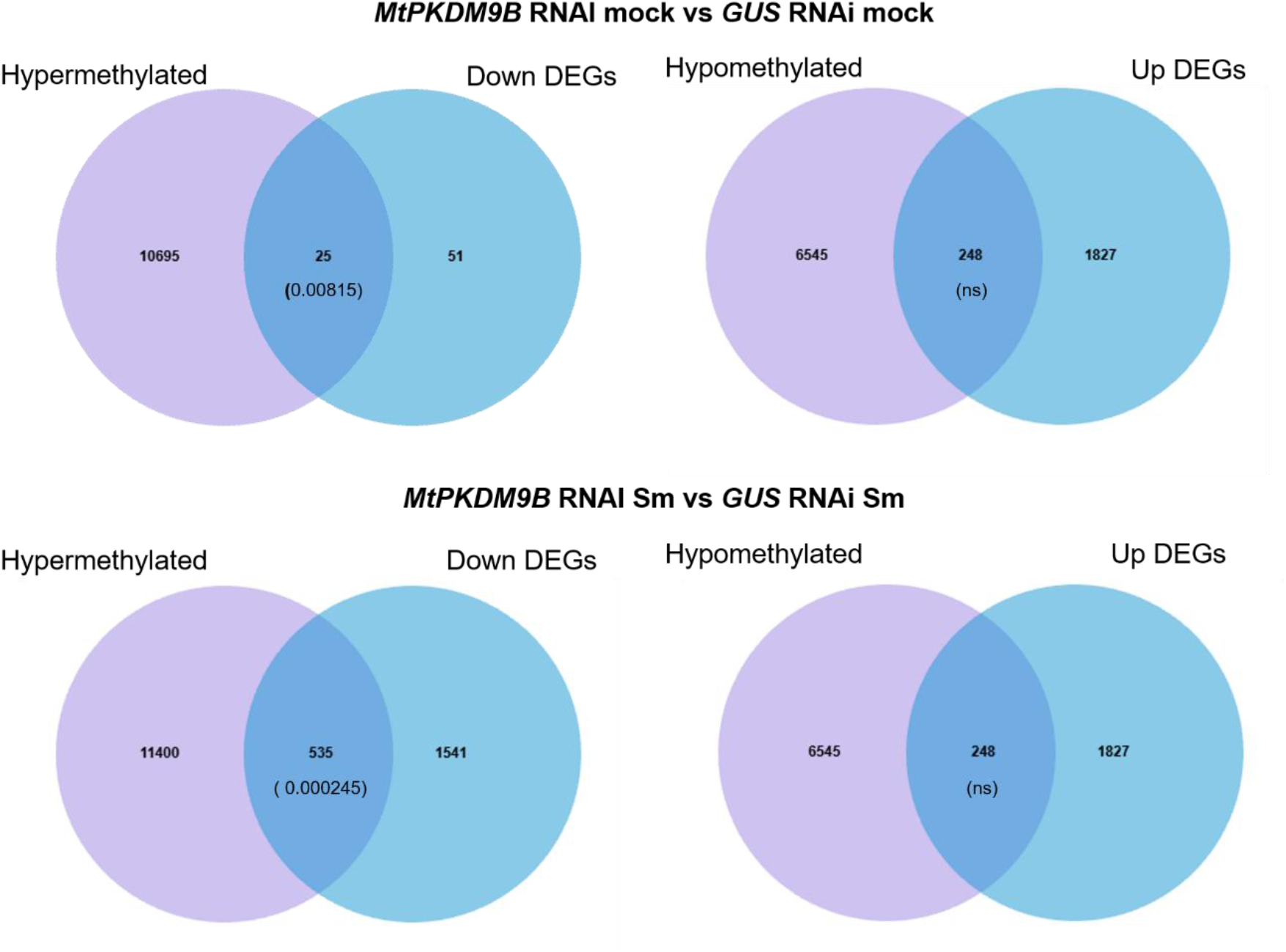
Genes with changes in H3K27me3 levels and DEGs between *GUS* RNAi and *MtPKDM9B* RNAi roots. Venn diagrams showing overlap between genes with hypermethylated H3K27me3 levels and up or down regulated DEGs or hypomethylated H3K27me3 levels and up or down regulated DEGs between *GUS* RNAi and *MtPKDM9B* RNAi roots in mock (upper panel) and *S. meliloti* inoculated (Sm) roots (lower panel). p-value for Fisher’s exact test is shown in brackets

## Notes

### Competing Interest Statement

The authors have declared no competing interest.

## References

Andriankaja A, Boisson-Dernier Al, Frances L, Sauviac L, Jauneau A, Barker DG, de Carvalho-Niebel F (2007) AP2-ERF Transcription Factors Mediate Nod Factor–Dependent Mt ENOD11 Activation in Root Hairs via a Novel cis-Regulatory Motif. The Plant Cell 19: 2866–2885

Ane JM, Kiss GB, Riely BK, Penmetsa RV, Oldroyd GE, Ayax C, Levy J, Debelle F, Baek JM, Kalo P, Rosenberg C, Roe BA, Long SR, Denarie J, Cook DR (2004) Medicago truncatula DMI1 required for bacterial and fungal symbioses in legumes. Science 303: 1364–1367

Antunez-Sanchez J, Naish M, Ramirez-Prado JS, Ohno S, Huang Y, Dawson A, Opassathian K, Manza-Mianza D, Ariel F, Raynaud C, Wibowo A, Daron J, Ueda M, Latrasse D, Slotkin RK, Weigel D, Benhamed M, Gutierrez-Marcos J (2020) A new role for histone demethylases in the maintenance of plant genome integrity. eLife 9: e58533

Baudin M, Laloum T, Lepage A, Ripodas C, Ariel F, Frances L, Crespi M, Gamas P, Blanco FA, Zanetti ME, de Carvalho-Niebel F, Niebel A (2015) A Phylogenetically Conserved Group of Nuclear Factor-Y Transcription Factors Interact to Control Nodulation in Legumes. Plant Physiol 169: 2761–2773

Boisson-Dernier A, Chabaud M, Garcia F, Becard G, Rosenberg C, Barker DG (2001) Agrobacterium rhizogenes-transformed roots of Medicago truncatula for the study of nitrogen-fixing and endomycorrhizal symbiotic associations. Mol Plant Microbe Interact 14: 695–700

Breakspear A, Liu C, Roy S, Stacey N, Rogers C, Trick M, Morieri G, Mysore KS, Wen J, Oldroyd GE, Downie JA, Murray JD (2014) The root hair “infectome” of Medicago truncatula uncovers changes in cell cycle genes and reveals a requirement for Auxin signaling in rhizobial infection. Plant Cell 26: 4680–4701

Cattaneo P, Graeff M, Marhava P, Hardtke CS (2019) Conditional effects of the epigenetic regulator JUMONJI 14 in Arabidopsis root growth. Development 146

Cerri MR, Frances L, Kelner A, Fournier J, Middleton PH, Auriac MC, Mysore KS, Wen J, Erard M, Barker DG, Oldroyd GE, de Carvalho-Niebel F (2016) The Symbiosis-Related ERN Transcription Factors Act in Concert to Coordinate Rhizobial Host Root Infection. Plant Physiol 171: 1037–1054

Cerri MR, Frances L, Laloum T, Auriac MC, Niebel A, Oldroyd GE, Barker DG, Fournier J, de Carvalho-Niebel F (2012) Medicago truncatula ERN transcription factors: regulatory interplay with NSP1/NSP2 GRAS factors and expression dynamics throughout rhizobial infection. Plant Physiol 160: 2155–2172

Cervantes-Perez SA, Thibivilliers S, Laffont C, Farmer AD, Frugier F, Libault M (2022) Cell-specific pathways recruited for symbiotic nodulation in the Medicago truncatula legume. Mol Plant 15: 1868–1888

Charpentier M, Sun J, Vaz Martins T, Radhakrishnan GV, Findlay K, Soumpourou E, Thouin J, Very AA, Sanders D, Morris RJ, Oldroyd GE (2016) Nuclear-localized cyclic nucleotide-gated channels mediate symbiotic calcium oscillations. Science 352: 1102–1105

Cook D, Dreyer D, Bonnet D, Howell M, Nony E, VandenBosch K (1995) Transient induction of a peroxidase gene in Medicago truncatula precedes infection by Rhizobium meliloti. Plant Cell 7: 43–55

Couzigou JM, Zhukov V, Mondy S, Abu el Heba G, Cosson V, Ellis TH, Ambrose M, Wen J, Tadege M, Tikhonovich I, Mysore KS, Putterill J, Hofer J, Borisov AY, Ratet P (2012) NODULE ROOT and COCHLEATA maintain nodule development and are legume orthologs of Arabidopsis BLADE-ON-PETIOLE genes. Plant Cell 24: 4498–4510

Crevillén P, Yang H, Cui X, Greeff C, Trick M, Qiu Q, Cao X, Dean C (2014) Epigenetic reprogramming that prevents transgenerational inheritance of the vernalized state. Nature 515: 587–590

Cui X, Lu F, Qiu Q, Zhou B, Gu L, Zhang S, Kang Y, Cui X, Ma X, Yao Q, Ma J, Zhang X, Cao X (2016) REF6 recognizes a specific DNA sequence to demethylate H3K27me3 and regulate organ boundary formation in Arabidopsis. Nat Genet 48: 694–699

de Lucas M, Pu L, Turco G, Gaudinier A, Morao AK, Harashima H, Kim D, Ron M, Sugimoto K, Roudier F, Brady SM (2016) Transcriptional Regulation of Arabidopsis Polycomb Repressive Complex 2 Coordinates Cell-Type Proliferation and Differentiation. The Plant Cell 28: 2616–2631

Dong W, Zhu Y, Chang H, Wang C, Yang J, Shi J, Gao J, Yang W, Lan L, Wang Y, Zhang X, Dai H, Miao Y, Xu L, He Z, Song C, Wu S, Wang D, Yu N, Wang E (2021) An SHR-SCR module specifies legume cortical cell fate to enable nodulation. Nature 589: 586–590

Fotouhi O, Nizamuddin S, Falk S, Schilling O, Knüchel-Clarke R, Biniossek ML, Timmers HTM (2023) Alternative mRNA Splicing Controls the Functions of the Histone H3K27 Demethylase UTX/KDM6A. Cancers 15: 3117

Franssen HJ, Xiao TT, Kulikova O, Wan X, Bisseling T, Scheres B, Heidstra R (2015) Root developmental programs shape the Medicago truncatula nodule meristem. Development 142: 2941–2950

Gan E-S, Xu Y, Wong J-Y, Geraldine Goh J, Sun B, Wee W-Y, Huang J, Ito T (2014) Jumonji demethylases moderate precocious flowering at elevated temperature via regulation of FLC in Arabidopsis. Nature Communications 5: 5098

Gao J-P, Xia C, Chiu CH, Chen Q, Jiang S, Wu X, Liang W, Sun J, Jhu M-Y, Wen J, Wang E, Murray JD, Oldroyd GED (2026) An NSP2-MYB module orchestrates flavonoid biosynthesis and nodule symbiosis. Current Biology 36: 940–953.e945

Groth M, Takeda N, Perry J, Uchida H, Draxl S, Brachmann A, Sato S, Tabata S, Kawaguchi M, Wang TL, Parniske M (2010) NENA, a Lotus japonicus homolog of Sec13, is required for rhizodermal infection by arbuscular mycorrhiza fungi and rhizobia but dispensable for cortical endosymbiotic development. Plant Cell 22: 2509–2526

Gu X, Xu T, He Y (2014) A Histone H3 Lysine-27 Methyltransferase Complex Represses Lateral Root Formation in *Arabidopsis thaliana*. Molecular Plant 7: 977-988

Guefrachi I, Nagymihaly M, Pislariu CI, Van de Velde W, Ratet P, Mars M, Udvardi MK, Kondorosi E, Mergaert P, Alunni B (2014) Extreme specificity of NCR gene expression in Medicago truncatula. BMC Genomics 15: 712

Haider S, Farrona S (2024) Decoding histone 3 lysine methylation: Insights into seed germination and flowering. Curr Opin Plant Biol 81: 102598

Hobecker KV, Reynoso MA, Bustos-Sanmamed P, Wen J, Mysore KS, Crespi M, Blanco FA, Zanetti ME (2017) The MicroRNA390/TAS3 Pathway Mediates Symbiotic Nodulation and Lateral Root Growth. Plant Physiology 174: 2469–2486

Jacob Y, Stroud H, LeBlanc C, Feng S, Zhuo L, Caro E, Hassel C, Gutierrez C, Michaels SD, Jacobsen SE (2010) Regulation of heterochromatic DNA replication by histone H3 lysine 27 methyltransferases. Nature 466: 987–991

Jardinaud M-F, Fromentin J, Auriac M-C, Moreau S, Pecrix Y, Taconnat L, Cottret L, Aubert G, Balzergue S, Burstin J, Carrere S, Gamas P (2022) MtEFD and MtEFD2: Two transcription factors with distinct neofunctionalization in symbiotic nodule development. Plant Physiology 189: 1587–1607

Jhu M-Y, Oldroyd GED (2023) Dancing to a different tune, can we switch from chemical to biological nitrogen fixation for sustainable food security? PLOS Biology 21: e3001982

Karimi M, Depicker A, Hilson P (2007) Recombinational cloning with plant gateway vectors. Plant Physiol 145: 1144–1154

Karimi M, Inze D, Depicker A (2002) GATEWAY vectors for Agrobacterium-mediated plant transformation. Trends Plant Sci 7: 193–195

Kim D, Paggi JM, Park C, Bennett C, Salzberg SL (2019) Graph-based genome alignment and genotyping with HISAT2 and HISAT-genotype. Nat Biotechnol 37: 907–915

Kirolinko C, Hobecker K, Cueva M, Botto F, Christ A, Niebel A, Ariel F, Blanco FA, Crespi M, Zanetti ME (2024) A lateral organ boundaries domain transcription factor acts downstream of the auxin response factor 2 to control nodulation and root architecture in Medicago truncatula. New Phytol

Kirolinko C, Hobecker K, Wen J, Mysore KS, Niebel A, Blanco FA, Zanetti ME (2021) Auxin Response Factor 2 (ARF2), ARF3, and ARF4 Mediate Both Lateral Root and Nitrogen Fixing Nodule Development in Medicago truncatula. Front Plant Sci 12: 659061

Langmead B (2010) Aligning short sequencing reads with Bowtie. Curr Protoc Bioinformatics Chapter 11: Unit 11 17

Li Q, Li J, Tang X, Chu C, Wang J, Zhou D-X, Zhao Y (2026) PLETHORA transcription factors orchestrate epigenetic silencing of bivalent chromatin to promote root meristem development in rice. Molecular Plant 19: 134–150

Limpens E, Ovchinnikova E, Journet EP, Chabaud M, Cosson V, Ratet P, Duc G, Fedorova E, Liu W, Op den Camp R, Zhukov V, Tikhonovich I, Borisov A, Bisseling T (2011) IPD3 controls the formation of nitrogen-fixing symbiosomes in pea and Medicago Spp. Mol Plant Microbe Interact 24: 1333–1344

Liu C, Lu F, Cui X, Cao X (2010) Histone methylation in higher plants. Annu Rev Plant Biol 61: 395–420

Liu J, Deng J, Zhu F, Li Y, Lu Z, Qin P, Wang T, Dong J (2018) The MtDMI2-MtPUB2 Negative Feedback Loop Plays a Role in Nodulation Homeostasis. Plant Physiol 176: 3003–3026

Lopez L, Perrella G, Calderini O, Porceddu A, Panara F (2022) Genome-Wide Identification of Histone Modification Gene Families in the Model Legume Medicago truncatula and Their Expression Analysis in Nodules. Plants (Basel) 11

Love MI, Huber W, Anders S (2014) Moderated estimation of fold change and dispersion for RNA-seq data with DESeq2. Genome Biol 15: 550

Lu F, Cui X, Zhang S, Jenuwein T, Cao X (2011) Arabidopsis REF6 is a histone H3 lysine 27 demethylase. Nature Genetics 43: 715–719

Luo Z, Lin J-s, Zhu Y, Fu M, Li X, Xie F (2021) NLP1 reciprocally regulates nitrate inhibition of nodulation through SUNN-CRA2 signaling in Medicago truncatula. Plant Communications 2: 100183

Luo Z, Wang J, Li F, Lu Y, Fang Z, Fu M, Mysore KS, Wen J, Gong J, Murray JD, Xie F (2023) The small peptide CEP1 and the NIN-like protein NLP1 regulate NRT2.1 to mediate root nodule formation across nitrate concentrations. Plant Cell 35: 776–794

Masubelele NH, Dewitte W, Menges M, Maughan S, Collins C, Huntley R, Nieuwland J, Scofield S, Murray JA (2005) D-type cyclins activate division in the root apex to promote seed germination in Arabidopsis. Proc Natl Acad Sci U S A 102: 15694–15699

Meade HM, and Signer, E. R. (1977) Genetic mapping of Rhizobium meliloti. . Proc Natl Acad Sci U S A 74: 2076–2078

Mergaert P, Uchiumi T, Alunni B, Evanno G, Cheron A, Catrice O, Mausset AE, Barloy-Hubler F, Galibert F, Kondorosi A, Kondorosi E (2006) Eukaryotic control on bacterial cell cycle and differentiation in the Rhizobium-legume symbiosis. Proc Natl Acad Sci U S A 103: 5230–5235

Millan-Zambrano G, Burton A, Bannister AJ, Schneider R (2022) Histone post-translational modifications - cause and consequence of genome function. Nat Rev Genet 23: 563–580

Morère-Le Paven M-C, Clochard T, Limami AM (2024) NPF and NRT2 from Pisum sativum Potentially Involved in Nodule Functioning: Lessons from Medicago truncatula and Lotus japonicus. Plants 13: 322

Morgan MAJ, Shilatifard A (2020) Reevaluating the roles of histone-modifying enzymes and their associated chromatin modifications in transcriptional regulation. Nat Genet 52: 1271–1281

Noh B, Lee S-H, Kim H-J, Yi G, Shin E-A, Lee M, Jung K-J, Doyle MR, Amasino RM, Noh Y-S (2004) Divergent Roles of a Pair of Homologous Jumonji/Zinc-Finger–Class Transcription Factor Proteins in the Regulation of Arabidopsis Flowering Time. The Plant Cell 16: 2601–2613

Noh B, Lee SH, Kim HJ, Yi G, Shin EA, Lee M, Jung KJ, Doyle MR, Amasino RM, Noh YS (2004) Divergent roles of a pair of homologous jumonji/zinc-finger-class transcription factor proteins in the regulation of Arabidopsis flowering time. Plant Cell 16: 2601–2613

Osipova MA, Mortier V, Demchenko KN, Tsyganov VE, Tikhonovich IA, Lutova LA, Dolgikh EA, Goormachtig S (2012) Wuschel-related homeobox5 gene expression and interaction of CLE peptides with components of the systemic control add two pieces to the puzzle of autoregulation of nodulation. Plant Physiol 158: 1329–1341

Pecrix Y, Staton SE, Sallet E, Lelandais-Briere C, Moreau S, Carrere S, Blein T, Jardinaud MF, Latrasse D, Zouine M, Zahm M, Kreplak J, Mayjonade B, Satge C, Perez M, Cauet S, Marande W, Chantry-Darmon C, Lopez-Roques C, Bouchez O, Berard A, Debelle F, Munos S, Bendahmane A, Berges H, Niebel A, Buitink J, Frugier F, Benhamed M, Crespi M, Gouzy J, Gamas P (2018) Whole-genome landscape of Medicago truncatula symbiotic genes. Nat Plants 4: 1017–1025

Peiter E, Sun J, Heckmann AB, Venkateshwaran M, Riely BK, Otegui MS, Edwards A, Freshour G, Hahn MG, Cook DR, Sanders D, Oldroyd GE, Downie JA, Ane JM (2007) The Medicago truncatula DMI1 protein modulates cytosolic calcium signaling. Plant Physiol 145: 192–203

Penmetsa RV, Cook DR (1997) A Legume Ethylene-Insensitive Mutant Hyperinfected by Its Rhizobial Symbiont. Science 275: 527–530

Qian S, Wang Y, Ma H, Zhang L (2015) Expansion and Functional Divergence of Jumonji C-Containing Histone Demethylases: Significance of Duplications in Ancestral Angiosperms and Vertebrates. Plant Physiol 168: 1321–1337

Quandt HJ, Pühler, A., and Broer, I. (1993) Transgenic root nodules of Vicia hirsuta: A fast and efficient system for the study of gene expression in indeterminate-type nodules. Molecular Plant-Microbe Interactions: 699–706

Ramírez F, Ryan DP, Grüning B, Bhardwaj V, Kilpert F, Richter AS, Heyne S, Dündar F, Manke T (2016) deepTools2: a next generation web server for deep-sequencing data analysis. Nucleic Acids Res 44: W160–165

Reynoso MA, Blanco FA, Bailey-Serres J, Crespi M, Zanetti ME (2013) Selective recruitment of mRNAs and miRNAs to polyribosomes in response to rhizobia infection in Medicago truncatula. Plant J 73 289–301

Sainz MM, Filippi CV, Eastman G, Sotelo-Silveira M, Martinez CM, Borsani O, Sotelo-Silveira J (2022) Polysome Purification from Soybean Symbiotic Nodules. J Vis Exp

Schiessl K, Lilley JLS, Lee T, Tamvakis I, Kohlen W, Bailey PC, Thomas A, Luptak J, Ramakrishnan K, Carpenter MD, Mysore KS, Wen J, Ahnert S, Grieneisen VA, Oldroyd GED (2019) NODULE INCEPTION Recruits the Lateral Root Developmental Program for Symbiotic Nodule Organogenesis in Medicago truncatula. Curr Biol 29: 3657–3668 e3655

Schwab R, Palatnik JF, Riester M, Schommer C, Schmid M, Weigel D (2005) Specific effects of microRNAs on the plant transcriptome. Dev Cell 8: 517–527

Shen L, Shao NY, Liu X, Maze I, Feng J, Nestler EJ (2013) diffReps: detecting differential chromatin modification sites from ChIP-seq data with biological replicates. PLoS One 8: e65598

Smit P, Raedts J, Portyanko V, Debelle F, Gough C, Bisseling T, Geurts R (2005) NSP1 of the GRAS protein family is essential for rhizobial Nod factor-induced transcription. Science 308: 1789–1791

Soyano T, Shimoda Y, Kawaguchi M, Hayashi M (2019) A shared gene drives lateral root development and root nodule symbiosis pathways in *Lotus*. Science 366: 1021–1023

Tarayre S, Vinardell JM, Cebolla A, Kondorosi A, Kondorosi E (2004) Two classes of the CDh1-type activators of the anaphase-promoting complex in plants: novel functional domains and distinct regulation. Plant Cell 16: 422–434

Tarayre S, Vinardell JM, Cebolla A, Kondorosi A, Kondorosi E (2004) Two Classes of the Cdh1-Type Activators of the Anaphase-Promoting Complex in Plants: Novel Functional Domains and Distinct Regulation[W]. The Plant Cell 16: 422–434

Thorvaldsdottir H, Robinson JT, Mesirov JP (2013) Integrative Genomics Viewer (IGV): high-performance genomics data visualization and exploration. Brief Bioinform 14: 178–192

Tian CF, Garnerone AM, Mathieu-Demaziere C, Masson-Boivin C, Batut J (2012) Plant-activated bacterial receptor adenylate cyclases modulate epidermal infection in the Sinorhizobium meliloti-Medicago symbiosis. Proc Natl Acad Sci U S A 109: 6751–6756

Traubenik S, Reynoso MA, Hobecker K, Lancia M, Hummel M, Rosen B, Town C, Bailey-Serres J, Blanco F, Zanetti ME (2020) Reprogramming of Root Cells during Nitrogen-Fixing Symbiosis Involves Dynamic Polysome Association of Coding and Noncoding RNAs. The Plant Cell 32: 352–373

Tu T, Gao Z, Li L, Chen J, Ye K, Xu T, Mai S, Han Q, Chen C, Wu S, Dong Y, Chen J, Huang L, Guan Y, Xie F, Chen X (2024) Soybean symbiotic-nodule zonation and cell differentiation are defined by NIN2 signaling and GH3-dependent auxin homeostasis. Developmental Cell 59: 2254–2269.e2256

Van de Velde W, Zehirov G, Szatmari A, Debreczeny M, Ishihara H, Kevei Z, Farkas A, Mikulass K, Nagy A, Tiricz H, Satiat-Jeunemaître B, Alunni B, Bourge M, Kucho K, Abe M, Kereszt A, Maroti G, Uchiumi T, Kondorosi E, Mergaert P (2010) Plant peptides govern terminal differentiation of bacteria in symbiosis. Science 327: 1122–1126

Wang F, Zhang L, Xie W, Cai X, Wang J, Wu X, Wang Y, Dong A, Su W (2025) ELF6 mutation suppresses the dwarf phenotype of Arabidopsis ino80 mutant by modulating cell cycle progression. New Phytol 248: 231–249

Wang X, Gao J, Gao S, Song Y, Yang Z, Kuai B (2019) The H3K27me3 demethylase REF6 promotes leaf senescence through directly activating major senescence regulatory and functional genes in Arabidopsis. PLoS Genet 15: e1008068

Xiao J, Lee U-S, Wagner D (2016) Tug of war: adding and removing histone lysine methylation in Arabidopsis. Current Opinion in Plant Biology 34: 41–53

Yan W, Chen D, Smaczniak C, Engelhorn J, Liu H, Yang W, Graf A, Carles CC, Zhou D-X, Kaufmann K (2018) Dynamic and spatial restriction of Polycomb activity by plant histone demethylases. Nature Plants 4: 681–689

Yan W, Chen D, Smaczniak C, Engelhorn J, Liu H, Yang W, Graf A, Carles CC, Zhou DX, Kaufmann K (2018) Dynamic and spatial restriction of Polycomb activity by plant histone demethylases. Nat Plants 4: 681–689

Yao ZL, Fang QF, Li JY, Zhou M, Du S, Chen HJ, Wang H, Jiang S-J, Wang X, Zhao Y, Ji XS (2023) Alternative splicing of histone demethylase *Kdm6bb* mediates temperature-induced sex reversal in the Nile tilapia. Current Biology 33: 5057–5070.e5055

Zanetti ME, Blanco F, Ferrari M, Ariel F, Benoit M, Niebel A, Crespi M (2024) Epigenetic control during root development and symbiosis. Plant Physiology 196: 697–710

Zibetti C, Adamo A, Binda C, Forneris F, Toffolo E, Verpelli C, Ginelli E, Mattevi A, Sala C, Battaglioli E (2010) Alternative splicing of the histone demethylase LSD1/KDM1 contributes to the modulation of neurite morphogenesis in the mammalian nervous system. J Neurosci 30: 2521–2532

## Supplemental references

Ge SX, Jung D, Yao R (2019) ShinyGO: a graphical gene-set enrichment tool for animals and plants. Bioinformatics 36: 2628–2629

Madeira F, Madhusoodanan N, Lee J, Eusebi A, Niewielska A, Tivey ARN, Lopez R, Butcher S (2024) The EMBL-EBI Job Dispatcher sequence analysis tools framework in 2024. Nucleic acids research 52: W521–W525

