## Supplemental Table 9 for "The plant specific Histone Lysine Demethylase MtPKDM9B mediates the root nodule symbiosis by controlling H3K27me3 levels and expression of symbiotic genes"

**Supplemental Table 9. Primers used in this study**

| Gene name | Gen ID | Primer | Sequence (5´-3´) | Reference |
| --- | --- | --- | --- | --- |
| *MtPKDM9B* | MtrunA17Chr1g0199511 | qMtPKDM9B F | CCTCGTGCTTCAGGCTCACT | Thiis work |
|  |  | qMtPKDM9B R | GCAGTTGCTGATGGGAGAGCA |  |
|  |  | MtPKDM9B.1 F | ATCACCTGAGGTAGTTGTTG |  |
|  |  | MtPKDM9B.1 R | GGCTGAATCCTACATGATAA |  |
|  |  | MtPKDM9B RNAi F | CACCATGTAGGATTCAGCCACGGT |  |
|  |  | MtPKDM9B RNAi R | ACGGTCTCTCAGACGAGAACTG |  |
| *MtPKDM9A* | MtrunA17Chr3g0116441 | MtPKDM9A F | CAACTCACACCACTCTGCTCA |  |
|  |  | MtPKDM9A R | GGTTGTTCTGTGCTGTGATTGTTG |  |
| *HISTONE LIKE 3 (HISL3)* | MtrunA17Chr4R0220620 | HISL3 F | ATTCCAAAGGCGGCTGCATA | Ariel et al., (2010) |
|  |  | HISL3 R | CTTTGCTTGGTGCTGTTTAGATGG |  |
| I miR-s | microRNA 319 | miR319 | GATGATTCTGTACTGACCTGCGATCTCTCTTTTGTATTCC | This work |
| II miR-a | microRNA 319 | miR319 | GATCGCAGGTCAGTACAGAATCATCAAAGAGAATCAATGA |  |
| III miR*s | microRNA 319 | miR319 | GATCACAGGTCAGTAGAGAATCTTCACAGGTCGTGATATG |  |
| IV miR*a | microRNA 319 | miR319 | GAAGATTCTCTACTGACCTGTGATCTACATATATATTCCT |  |
| A | microRNA 319 | miR319 | CTGCAAGGCGATTAAGTTGGG TAAC |  |
| B | microRNA 319 | miR319 | GCGGATAACAATTTCACACAG GAAACAG |  |
| *MtrunA17Chr3R0162790* | *MtrunA17Chr3R0162790* | *MtrunA17Chr3R0162790 F* | GGAGTAGGCACATACCCTGC |  |
|  |  | *MtrunA17Chr3R0162790 R* | GCAACTTCTGAGCCCATAGTCAT |  |
| *MtACTIN11* | MtrunA17Chr7g0223901 | Act11 F | ACCCAAAGCATCAAATAATAAGTCAACC | Ariel et al., (2010) |
|  |  | Act11 R | ACCCAAAGCATCAAATAATAAGTCAACC |  |
| *MtERN1* | MtrunA17Chr7g0253424 | ERN1 F | GGAAGATGGTGCTGTTGCTT | Andriankaja et al., (2007) |
|  |  | ERN1 R | TGTTGGATTGTGAACCTGACTC |  |
| *MtLBD16* | MtrunA17Chr7g0260971 | ChIP LBD16 F | AGTCCCTAATTGCACCTGC | This work |
|  |  | ChIP LBD16 R | ATTGTTGCTAGGCACACGGA |  |
| *MtSTYL1* | MtrunA17Chr1g0155791 | ChIP STYL1 F | GAGGAGGTTGTGGTACTG |  |
|  |  | ChIP STYL1 R | GGTCAAATTCCGTGTCTC | Schiessl et al., (2019) |
|  |  | qMtSTYL1 F | AGCAGCAGCAACAACAGTTTCAC |  |
|  |  | qSTYL1 R | AAATTTCCCAACTCCAACCCTGTG |  |
| *MtNSP1* | MtrunA17Chr8g0344101 | ChIP NSP1 F | CACATGGTTTGAGTGAGTT | This work |
|  |  | ChIP NSP1 R | TATTTTGCCACACCAATATC |  |
|  |  | qNSP1 F | GTGGTTAGAAAATCTGGTGGG | Smit et al., (2005) |
|  |  | qNSP1 R | GTGTCAATGCTCGAAGACCA |  |

**References**

**Ariel FD, Diet A, Crespi M, Chan RL** (2010) The LOB-like transcription factor Mt LBD1 controls Medicago truncatula root architecture under salt stress. Plant signaling & behavior **5**: 1666–1668.

**Andriankaja A, Boisson-Dernier A, Frances L, Sauviac L, Jauneau A, Barker DG, de Carvalho-Niebel F** (2007) AP2-ERF Transcription Factors Mediate Nod Factor Dependent Mt ENOD11 Activation in Root Hairs via a Novel cis-Regulatory Motif. THE PLANT CELL ONLINE. doi: 10.1105/tpc.107.052944

**Schiessl K, Lilley JLS, Lee T, Tamvakis I, Kohlen W, Bailey PC, Thomas A, Luptak J, Ramakrishnan K, Carpenter MD, et al** (2019) NODULE INCEPTION Recruits the Lateral Root Developmental Program for Symbiotic Nodule Organogenesis in Medicago truncatula. Current Biology **29**: 3657-3668.e5

**Smit P, Raedts J, Portyanko V, Debellé F, Gough C, Bisseling T, Geurts R** (2005) NSP1 of the GRAS protein family is essential for rhizobial Nod factor-induced transcription. Science (New York, NY) **308**: 1789–1791
